# Sublayer- and cell-type-specific neurodegenerative transcriptional trajectories in hippocampal sclerosis

**DOI:** 10.1101/2021.02.03.429560

**Authors:** Elena Cid, Angel Marquez-Galera, Manuel Valero, Beatriz Gal, Daniel C. Medeiros, Carmen M. Navarron, Luis Ballesteros-Esteban, Rita Reig-Viader, Aixa V. Morales, Ivan Fernandez-Lamo, Daniel Gomez-Dominguez, Masaaki Sato, Yasunori Hayashi, Alex Bayes, Angel Barco, Jose P Lopez-Atalaya, Liset M de la Prida

## Abstract

Hippocampal sclerosis, the major neuropathological hallmark of temporal lobe epilepsy, is characterized by different patterns of neuronal loss. The mechanisms of cell-type specific vulnerability, their progression and histopathological classification remain controversial. Here using single-cell electrophysiology in vivo and immediate early gene expression, we reveal that superficial CA1 pyramidal neurons are overactive in epileptic rats and mice *in vivo*. Bulk tissue and single-nucleus expression profiling disclosed sublayer-specific transcriptomic signatures and robust microglial pro-inflammatory responses. Transcripts regulating neuronal processes such as voltage-channels, synaptic signalling and cell adhesion molecules were deregulated by epilepsy differently across sublayers, while neurodegenerative signatures primarily involved superficial cells. Pseudotime analysis of gene expression in single-nuclei and *in situ* validation revealed separated trajectories from health to epilepsy across cell types, and identified a subset of superficial cells undergoing a later stage in neurodegeneration. Our findings indicate sublayer- and cell type-specific changes associated with selective CA1 neuronal damage contributing to progression of hippocampal sclerosis.

## Introduction

Epilepsies are brain disorders characterized by enduring predisposition to generate seizures with emotional and cognitive associated comorbidities. Despite significant therapeutic advances, one-third of patients remain resistant to pharmacotherapy (*1*). Temporal lobe epilepsy (TLE), the most prevalent form of pharmacoresistant epilepsy, is frequently associated with hippocampal sclerosis (*2*). Hippocampal sclerosis is characterized by specific patterns of neuronal loss affecting different hippocampal subfields from CA1 to CA3/4 areas, the hilus of the dentate gyrus and superficial layers of the entorhinal cortex (*3–5*). Factors such as epilepsy history, age of onset and relationship with early precipitating events may all influence the degree and severity of hippocampal sclerosis (*6*).

The most common form of hippocampal sclerosis (type 1; 60-80% TLE cases) shows severe neuronal loss of CA1, CA3 and CA4 pyramidal neurons and milder loss in CA2, with variability along the anteroposterior axis (*2, 7*). Other cellular types, including microglia and astrocytes, are also affected (*2, 8*). In contrast, type 2 hippocampal sclerosis (10-20% cases) is associated with predominant CA1 neurodegeneration and minimal loss in other regions (*2*). Given disparities between clinical series, there is no consensus yet on whether neuronal loss progresses or not along years (*5, 6*). In addition, individual variabilities and anatomical inhomogeneities complicate classification (*9–11*). For instance, patchy neuronal loss has been described in the CA1 region in some cases, while in others it rather seems to adopt a more laminar profile (*12*). In some patients, cell loss concentrates in the CA4 region and the dentate gyrus, and is frequently integrated in dual pathologies, e.g. TLE and malformations of the cortical development, classified as type 3 hippocampal sclerosis (3-7%) (*13*). The mechanisms underlying specific vulnerability of diverse cells, their role in the histopathological landscape and clinical significance remain unknown.

Recent techniques and methods operating at single-cell resolution point to an exquisite cell-type specific organization that is instrumental for brain function (*14, 15*). In the hippocampus, the CA1 region is organized radially in two distinct sublayers with characteristic gene expression gradients along the anteroposterior and proximodistal axes (*16–18*). Functionally, superficial (closer to radiatum) and deep (closer to oriens) CA1 pyramidal neurons project differentially and diverge in their participation of sharp-wave ripple activity, theta-gamma oscillations and behavioural-cognitive correlates (*19, 20*). In spite of data suggesting critical regionalization of CA1 neuronal responses to ischemia, anoxia and epilepsy (*21–23*), little is known on their clinical relevance and potential relationship with neuronal vulnerability. Understanding the impact of cellular diversity and transcriptional changes in epilepsy progression may lend insights into more specific mechanisms towards new diagnostic and therapeutic opportunities.

Here, we combine gene expression profiling at the single-nucleus and microdissected tissue levels with single-cell electrophysiology to disclose epileptogenic and neurodegenerative changes running differentially across CA1 sublayers in an experimental model of hippocampal sclerosis. Our study highlights the importance of leveraging on cell type specificity to better understand the phenotypic complexities accompanying hippocampal sclerosis in epilepsy.

## Results

Animals used in this study were examined in the chronic phase of the *status epilepticus* model of TLE (6-12 weeks post-*status*) when they already exhibited spontaneous seizures and interictal discharges (IID) (Sup.Fig.1A,B). In epileptic rats and mice, these events were typically associated with high-frequency oscillations (HFOs; Sup.Fig. 1B), which are considered biomarkers of epileptogenesis (*23, 24*). We focused in the dorsal hippocampus given the major role in associated cognitive comorbidities of epilepsy and more consistent neuronal loss as compared to ventral (*23, 25*).

### Large activity burden in superficial CA1 pyramidal cells during epileptiform activities

Single CA1 pyramidal cells were intracellularly recorded from chronic epileptic rats to evaluate their intrinsic excitability and activity during IID, sharp-wave (SPW) fast ripples and ictal discharges detected with multi-site silicon probes under urethane anesthesia (Fig.1A-D). Recorded cells were identified post hoc (with streptavidin) and immunostained against Calbindin, to classify them as deep (negative) or superficial pyramidal cells (positive) (Fig.1A). SPW-associated HFO events were automatically detected and classified as ripples (100-150 Hz), fast ripples (>150 Hz) and IID using amplitude and spectral information (Sup.Fig.1C, D), as before (*23*).

**Fig.1.**
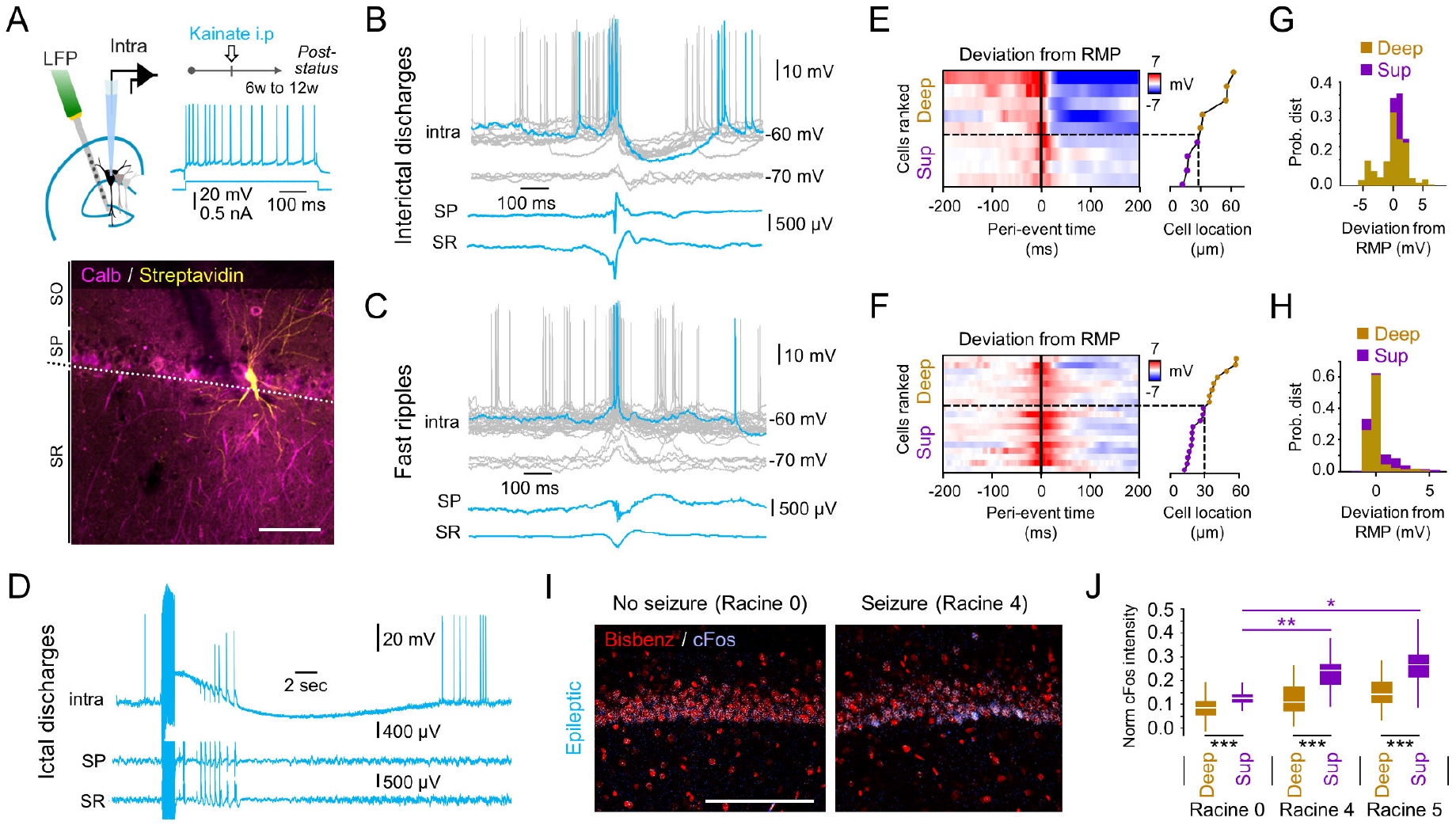
Differential responses of CA1 pyramidal cells during epileptiform activities. **A**, Intracellular and multi-site LFP recordings were obtained from urethane anesthetized epileptic rats. Cells were identified with streptavidin and tested against Calbindin. Scale bar 50 μm. **B**, Intracellular activity during IID at different membrane potentials (gray traces) in the representative cell shown in A. Traces are aligned by the peak of IID recorded at the stratum radiatum (SR). HFOs were recorded at the stratum pyramidale (SP). **C**, Responses of the cell shown in B during sharp-wave fast ripples. **D**, Intracellular ictal discharges associated to a non-convulsive seizure evoked by electrical stimulation of Schaffer collaterals. **E**, Deviation from the resting membrane potential (RMP) recorded in individual cells during interictal discharges. Red colors reflect depolarization, blue hyperpolarization. Cells are ranked by their distance to SR and classified as deep and superficial (subplot at right). The discontinuous line marks sublayer limits. **F**, Same as in E for SPW-fast ripples. **G**, Mean membrane potential responses around IID events showed differences between deep and superficial cells (Friedman Chi2(1,333)=46.2049, p<0.001). Data from n=4 deep and n=5 superficial CA1 pyramidal cells. **H**, Same as in G for SPW-fast ripple events. Note larger after-event depolarization in superficial cells (Friedman Chi2(1,333)=14.67, p<0.0001). Data from n=8 deep and n=11 superficial CA1 pyramidal cells. **I**, c-Fos immunoreactivity detected 1h after sound stimulation in representative sections from one rat experiencing no seizure (Racine 0) and one rat exhibiting bilateral forelimb clonus with rearing (Racine 4). Scale 100 μm. **J**, Intensity of c-Fos from all pyramidal cells in one section per rat as a function of induced seizure severity (Racine scale). 2-way Friedman test effects for seizure severity (p<0.0001) and sublayer (p<0.0001). Posthoc mean rank differences ***, p<0.001. *, p<0.05; **, p<0.001.

Intracellular activities recorded during either IID events (n=9 cells; Fig.1B) and SPW-fast ripples (n=19 cells; Fig.1C) were typically associated with consistent depolarization from the resting membrane potential (RMP) and firing of all pyramidal cells examined (Fig.1E,F). A temporal analysis of membrane potential changes (30 ms bins) showed differences between cell types during both IID events (Fig.1G; Friedman Chi2(1,333)=3.88, p<0.001) and SPW-fast ripples (Fig.1H; Chi2(1,333)=14.67, p-value<0.0001), with superficial cells consistently showing larger depolarization. Post-event membrane potential responses recorded showed effect for the type of events (F(2,54)=5.95, p=0.0048), sublayer (F(1,54)=5.76, p=0.0202) and interaction (F(2,54)=4.74, p=0.0131). Post hoc Tuckey tests confirmed significant smaller hyperpolarization (p=0.0028; unpaired t-test) and higher firing rate (p=0.040) following IID events in superficial as compared with deep cells. We found some minor differences of intrinsic excitability between cell-types and groups (Sup.Table.1).

We reasoned that superficial pyramidal cells should be more steadily activated during seizures, given their poor post-event hyperpolarization. To evaluate this point we induced ictal discharges by repetitive 100 Hz stimulation (300 ms) of the contralateral CA3 region while recording from single cells. Cells recorded during ictal events exhibited variable long-lasting depolarizing shifts (3.8-13.5 sec; n=12 cells; Fig.1D) and only a minority were successfully recovered for histological validation (n=4 deep cells), preventing subsequent comparisons across sublayers. To circumvent this problem, we exposed 3 epileptic rats to high-pitched sounds to promote convulsive seizures (random pulses of 95-100 dB and 1-20 sec duration at 0.05-1 Hz during 10 min). Two rats experienced convulsive seizures with forelimb clonus and one rat remained unaffected, as judged by clinical criteria (Racine scale). Animals were sacrificed after 1 hour to evaluate expression of the immediate-early gene c-Fos as a function of seizure severity (Fig.1I). We noticed preferential expression of c-Fos in superficial pyramidal cells and significant effect of seizure severity (p<0.0001) and sublayer (p<0.0001; 2-way Friedman test) (Fig.1J).

Altogether, these results suggest higher responsiveness of superficial CA1 pyramidal cells during epileptiform activities in chronic epileptic rats.

### Sublayer regionalization of epilepsy-associated transcriptional responses

To investigate how the differential responsiveness of deep and superficial cells may relate with distinct transcriptional responses across sublayers, we performed RNA sequencing (RNAseq) analysis of laser-microdissected samples from control (saline-injected) and epileptic rats (n=3 replicates each; Fig.2A; Suppl.Fig.2A,B). A public application provides easy visualization of these data (http://lopezatalayalab.in.umh-csic.es/CA1_Sublayers_&_Epilepsy/) (see also Sup.Table 2).

**Fig.2.**
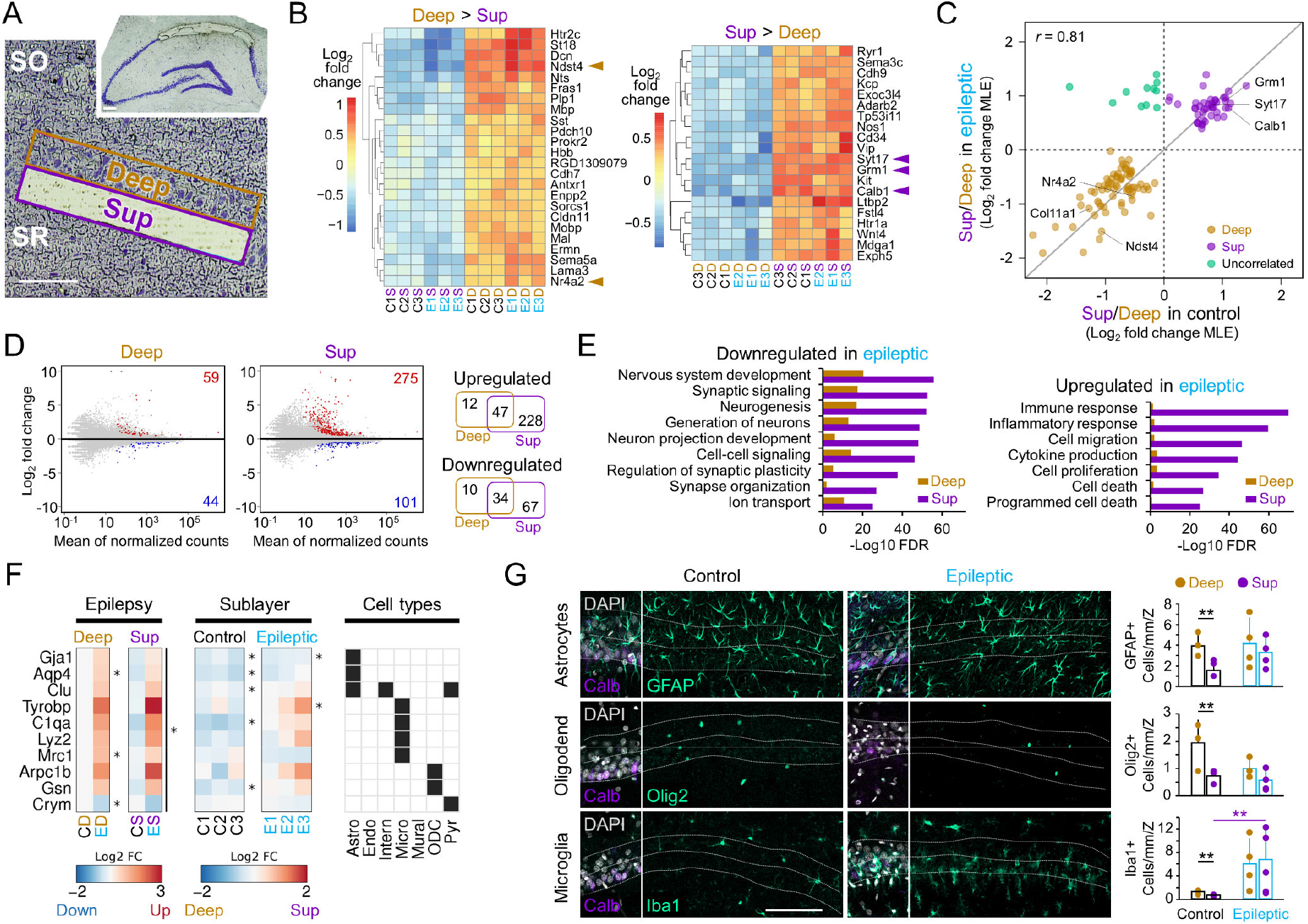
Bulk gene expression profiling of the epileptic hippocampal area CA1 reveals regionalized transcriptional responses. **A**, Representative LCM sampling of the superficial CA1 sublayer. Scale bar 100 μm. **B**, Heatmaps of genes differentially enriched in deep (D; left) and superficial CA1 sublayers (S; right) in three replicates from control (C) and epileptic (E) rats (Adj p-value<0.01 and LFC>0.5). Arrowheads point to *bona fide* gene markers of superficial (*Calb1, Grm1, Syt17*) and deep (*Ndst4*) pyramidal neurons. Note also the presence of gene markers for interneurons (e.g. *Vip, Sst, Sema3c, Kit*), and oligodendrocytes (*Plp1, Mbp, Mobp, Mal, Enpp2, Cldn11, Ermn*). **C**, Scatter plot of genes differentially expressed between superficial and deep CA1 sublayers in control or epileptic rats (Adj p-value<0.01). Note a subset of uncorrelated transcripts at the superficial sublayer in epileptic rats (green). MLE: maximum-likelihood estimate. **D**, MA plots showing epilepsy-associated DEGs in the deep and superficial sublayers (Adj p-value<0.01 and LFC>0.5). Venn diagrams at right show up- (red dots at left) and donwregulated DEGs (blue dots). **E**, GO analysis of upregulated and downregulated genes in the epileptic condition for deep and superficial sublayers (Adj p-value<0.01 and LFC>0.5). **F**, *Left:* Heatmaps showing mean differential expression (LFC) of cell markers genes in response to epilepsy (Epilepsy) in deep and superficial sublayers. *Middle:* Same as before but for sublayer effect (Sublayer) in control (Control) and epileptic (Epileptic) groups, shown as individual data. *Right:* Matrix shows cell types (Cell types) associated to each marker gene: Astro, astrocytes; Endo, endothelial cells; Intern, interneurons; Micro, microglia; Mural, mural cells; ODC, oligodendrocytes; Pyr, pyramidal cells. *, Adj p-value<0.1. **G**, *Left:* Representative control and epileptic sections immunostained against GFAP (Astrocytes; top), Olig2 (Oligodendrocytes; middle), and Iba1 (Microglia; bottom). The leftmost section on each row shows co-localization between the specific marker, Calb and DAPI. *Right:* Quantification of astrocyte (GFAP+), oligodendrocyte (Olig2+), and microglia (Iba1+) linear cell density across sublayers and groups. Significant differences of interaction between sublayer and groups for astrocytes (F(1,5)=10.1, p=0.022) and microglia (F(1,5)=7.4, p=0.042) with post hoc differences at deep-superficial layers in the control group. Effect of sublayer for oligodendrocytes (F(1,5)=23.8, p=0.0045. Differences between groups only for microglia cells. Data from n=3 control, n=4 epileptic rats. *, p<0.05; **, p<0.01 post hoc tests.

Analysis of LCM-RNAseq data revealed sublayer-specific genes common to control and epileptic animals (Fig.2B; deep versus superficial samples). Among these, we retrieved *bona fide* marker genes of superficial (*Calb1, Grm1*, and *Syt17*) and deep CA1 pyramidal cells (*Ndst4*), consistent with previous data in mice (*17, 26*) (Suppl.Fig.2C; see Suppl.Fig.2D,E for validation by in situ hybridization). Sublayer gene expression analysis confirmed preserved regionalization in epileptic rats with only a subset of uncorrelated transcripts (Fig.2C; green dots; see Suppl.Fig.2F for less stringent criteria: Adj p-value<0.1). Strikingly, we found that some gene markers of other cell types, including interneurons (*Sst, Vip, Kit*), oligodendrocytes (*Mbp, Plp1, Mobp*) and microglia (*Csf1r, Tgfbr1*) exhibited sublayer differences, consistent with heterogeneous distribution of cell types across the CA1 radial axis (Fig.2B; Suppl.Fig.2E) (see next section).

Next, we investigated the transcriptional changes associated to epilepsy across CA1 sublayers (control versus epileptic samples per sublayer). We identified 103 differentially expressed genes (DEGs) in the deep sublayer of control versus epileptic and 376 DEGs in the superficial sublayer (Fig.2D). While epilepsy-associated transcriptional changes run in similar direction in both sublayers (i.e. no counter-regulated genes), they were more exacerbated in the superficial CA1 (Fig.2D; Suppl.Fig.2H). Overall, we retrieved more upregulated than downregulated genes, particularly in the superficial (73% DEGs) versus the deep sublayer (57%) (Fig.2D; Venn diagrams at right). Functional enrichment analyses also revealed marked differences (Fig.2E). Downregulated DEGs typically involved Gene Ontology (GO) terms associated with neuronal processes, such as synaptic signaling, neuron projection development, regulation of synaptic plasticity, and synapse organization, including many well-known modulators of epileptogenic process such as *Cacna1a, Cntnap2, Kcnb1, Kcnip4, Gabbr2, Gria1, Grin2a and Nav2* amongst others (*27*) (Supp.Table.2). In contrast, significantly upregulated DEGs were more linked to immune and inflammatory responses, including cytokine production, cell migration, cell proliferation, and programmed cell death (e.g. *Cx3cr1, P2ry12, Tgfb1, Itgb1, Ripk3*) (Supp.Table.2). Most of these GO families were significantly enriched in the superficial CA1 sublayer (Fig.2E). Thus, differential gene expression analysis suggests that the nature and severity of transcriptional responses to TLE may segregate radially across CA1.

### Cell-type deconvolution of LCM-RNAseq data suggests sublayer-specific neurodegeneration associated to microglia

Given sublayer heterogeneity of biological processes revealed by bulk LCM-RNAseq, we reasoned that they might reflect different contribution by discrete cellular populations within the CA1 sample and/or changes of cell-type composition in response to epilepsy.

To address this point, we leveraged on a published dataset of *bona fide* transcripts from 3,005 barcoded individual cells from the mouse somatosensory S1 cortex and CA1 regions (*14*), (Suppl.Fig.3A). We performed unsupervised clustering to identify cell-type specific genes in LCM-RNAseq data, which were grouped in seven major classes: pyramidal cells, interneurons, astrocytes, mural and endothelial cells, microglia, and oligodendrocytes (Suppl.Fig.3B,C). In control animals, signatures of all these cell types were prominent in the CA1 deep layer, while genes enriched in the superficial sublayer were mostly associated to pyramidal neurons and interneurons (Suppl.Fig.3D,E). In contrast, we noticed strong upregulation of gene markers for microglia (Micro; e.g *Tyrobp*) and oligodendrocytes (ODC; e.g *Arpc1b*) at the superficial layers of the epileptic CA1 (Fig.2F). To evaluate this *in situ*, we performed immunofluorescence staining against protein markers for microglia (Iba1), astrocytes (GFAP), and oligodendrocytes (Olig2) and quantified their density across sublayers (Fig.2G, left). We confirmed the presence of a physiological segregation of glial cell types across the CA1 sublayers (control) and changes in TLE, with between-groups differences reaching significance for microglia (Fig.2G, right).

Prompted by these results, we dissected the contribution of microglia-associated transcripts in our sample. We noticed that most of the uncorrelated transcripts in the comparison of sublayer-enriched genes across conditions were highly expressed in microglia (green dots in Fig.2C and Suppl.Fig.2F; see microglia-specific genes in Suppl.Fig.2G). Thus, we evaluated their potential functional effect by building the microglial sensome (*28*), which was profoundly upregulated in the superficial sublayer of epileptic rats (Suppl.Fig.4A). Moreover, evaluation of microglia-neuron interactions via ligand-receptor pairing confirmed sublayer-specific effects (Suppl.Fig.4B). For example, in the epileptic superficial CA1 subfield we found dysregulated expression of the transcript encoding the CD200 receptor, whose expression in neurons provides a “don’t eat me” signal that reduces microglial activation (*29*). Similarly, *Cx3cl1*, a neuronal chemokine that dampens microglia inflammatory response and neurotoxicity was also strongly downregulated whereas *C3* and *Csf1* transcripts were upregulated in superficial CA1 sublayer (*30, 31*). In contrast, the epileptic deep CA1 sublayer showed a much weaker microglia signature and only *Cx3cl1* was found downregulated (Supp.Fig.4B).

Our analyses reveal the heterogeneous distribution of resident cell types across the CA1 radial axis of the normal hippocampus. Strikingly, we identified critical changes in the epileptic hippocampus, including local accumulation of reactive microglia and changes of key modulators of the neuronal immune response specifically at the superficial sublayer.

### CA1 hippocampal sclerosis is degenerative and sublayer-specific

Given the results described above, we sought to evaluate the anatomical landscape of CA1 hippocampal sclerosis. Previously, we reported significant reduction of linear cell density concentrated at the CA1 region in the systemic kainate model of TLE in rats, more consistent with sclerosis type 2 (*32*). To evaluate CA1 sublayers precisely, we combined immunostaining against the CA1 marker Wfs1 with the superficial marker Calbindin (Fig.3A). Proximal, intermediate and distal CA1 segments were evaluated separately in tissue from 10 control and 10 epileptic rats (1 section per animal, from −3.2 to −4.8 mm from bregma).

**Fig.3.**
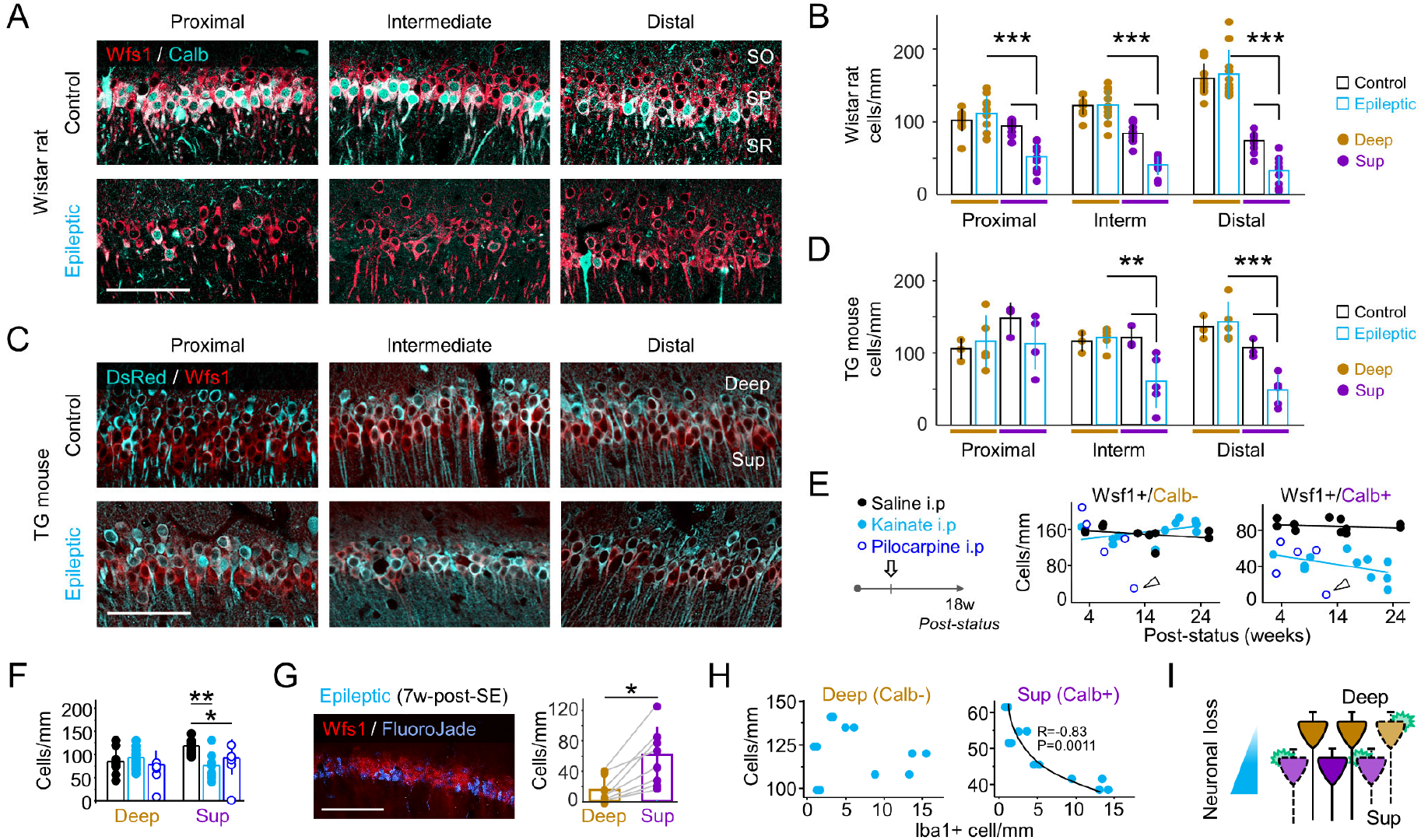
Hippocampal sclerosis is sublayer and cell-type specific. **A**, Immunostaining against the CA1 specific marker Wfs1 co-localized with Calb at proximal, intermediate and distal segments in control and epileptic rats. Scale bar, 100 μm. **B**, Quantification of CA1 linear cell density confirmed neuronal loss affecting mainly Calb+ cells. Data from n=10 control and n=10 kainate-treated epileptic rats (1 section per rat, −3.2 to −4.8 mm from bregma; 20x). Significant 3-way ANOVA for group (F1,120)=32.6, p<0.0001), proximodistal (F(2,120)=11.7, p<0.0001) and sublayer (F(1,120)=424,1; p<0.0001). Post hoc Tukey test: **, p<0.01; ***, p<0.005. **C**, Hippocampal sections from representative control and epileptic transgenic (TG) mouse expressing G-GaMP7-DsRed2 in Calb-deep pyramidal cells. **D**, Linear density data from n=3 control and n=5 epileptic TG mice (1 section per mouse at 20x). Significant 3-way ANOVA for groups (F(1,36)=7.8, p=0.008) and sublayers (F(1,36)=8.9, p=0.0051), no proximodistal effect. Post hoc t-test: **, p<0.01; ***, p<0.005. **E**, *Left:* Experimental timeline. *Right:* Mean density of Calb+ and Calb-CA1 pyramidal cells as a function of time. Data from n=10 control; n=10 epileptic rats (kainate) and n=5 epileptic rats (lithium-pilocarpine). Arrowheads point to a lithium-pilocarpine rat with full cell loss in CA1 10 weeks after status. **F**, CA1 mean cell loss in Wistar rats quantified as deep versus superficial. Superficial CA1 cells are more affected than deep cells. Significant effect for group in a 2-way ANOVA, F(2,48)=5.9, p=0.005. Post hoc unpaired t-test *, p<0.01, **, p<0.01. **G**, Fluoro-Jade signal co-localized with Wfs1 and quantification of Fluoro-Jade+ cells across sublayers; n=8 Wistar rats. Paired t-test *, p<0.01. **H**, Density of deep (Calb-; left) and superficial (Calb+; right) pyramidal cells against the density of microglia (Iba1+) counted at deep and superficial sublayers (data from n=6 epileptic rats). Significant correlation only for superficial. **I**, Schematic summary of histopathological findings.

Cell loss in all three segments was mostly restricted to Calb+ superficial CA1 pyramidal cells in chronic epileptic rats (Fig.3B; 3-way ANOVA for group, sublayer and proximo-distal effects all significant at p<0.0001; post hoc Tukey test p<0.001). Since Calbindin immunoreactivity may be affected during epileptogenesis (*33*), we exploited the transgenic mouse line Thy1.2-G-CaMP7-DsRed2 with restricted expression on deep pyramidal cells (Fig.3C). We found similar regionalization of neuronal loss at the intermediate and distal segments of chronic epileptic mice using co-localization between DsRed2 and Wfs1 (Fig.3D, left; n=3 control, n=5 epileptic; 6-8 weeks post-*status*; significant interaction between group and sublayer p=0.0006; post hoc t-test p<0.01). We also evaluated temporal trends of CA1 cell loss in epileptic rats where a time-dependent decrease of Calb+ cells was appreciated (r=-0.52; p=0.097; Pearson correlation), but not for control animals (p=0.66; Fig.3E) excluding age effects (*34*). We also examined sublayer-specific vulnerability of CA1 neurons in the lithium-pilocarpine model in rats (*35*), which is more consistent with type 1 hippocampal sclerosis (*32*). Notably, pilocarpine-treated rats showed similar reduction of Calb+ CA1 pyramidal cells early along epileptogenesis (p=0.0003; n=5 rats; 3-6 weeks post-*status*; Fig.3E, note full cell loss after 10 weeks post-*status*, arrowhead). Finally, to fully exclude interaction with Calbindin immunoreactivity, we evaluated Wfs1+ cell density at deep and superficial sublayers by relying only on anatomical criteria (location within sublayer) and found consistent results in both TLE models (Fig.3F; 2-way ANOVA for group p=0.005).

To further confirm sublayer-specific neurodegeneration, we combined Fluoro-Jade staining, which characteristically label degenerating cells, with Wfs1 immunostaining in sections from epileptic rats at different time points after kainate (2-23 weeks; n=6 epileptic and n=2 saline-injected rats) and again observed stronger neurodegeneration in superficial sublayers (Fig.3G; paired t-test p=0.0017). Notably, double immunostaining against Calb and the microglial marker Iba1 revealed an inversed correlation between the amount of microglia and the density of superficial pyramidal neurons (Calb+; r=-0.87, p=0.0236), but not for deep cells (Calb-; Fig.3H). This result indicates that loss of superficial pyramidal neurons is strongly associated to local accumulation of microglia.

Taken altogether, our results support the idea that hippocampal sclerosis results from specific interactions between different cell types in a particular niche (i.e. superficial CA1 pyramidal cells and activated microglia) leading to regionalized neurodegenerative signatures in the sclerotic CA1.

### Single nucleus RNAseq confirms cell-type specific neurodegeneration of CA1 pyramidal cells

To gain more insights into the transcriptional activity patterns underlying hippocampal sclerosis, we performed unbiased high-throughput RNA sequencing of isolated single-nuclei (snRNAseq) (Fig.4A). We analyzed the transcriptomes of 6,739 single high-quality CA1 barcoded nuclei derived from 2 control and 2 epileptic mice (n=3,661 nuclei, >710 unique molecular identifiers or UMI in control; n=3,078 nuclei, >540 UMI in epileptic samples). Consistent with recent reports (*14, 15*), barcoded cells were automatically classified in 13 clusters of CA1 nuclei that were then aggregated in six major cell classes: excitatory neurons, interneurons, oligodendrocytes, microglia, oligodendrocyte precursor cells and astrocytes (Suppl.Fig.5A,B; see Suppl.Fig5C for control versus epileptic and Suppl.Fig5D for UMI levels and number of genes per cell type) (Supp.Table 3). Data can be visualized in a public application (http://lopezatalayalab.in.umh-csic.es/CA1_SingleNuclei_&_Epilepsy/).

**Fig.4.**
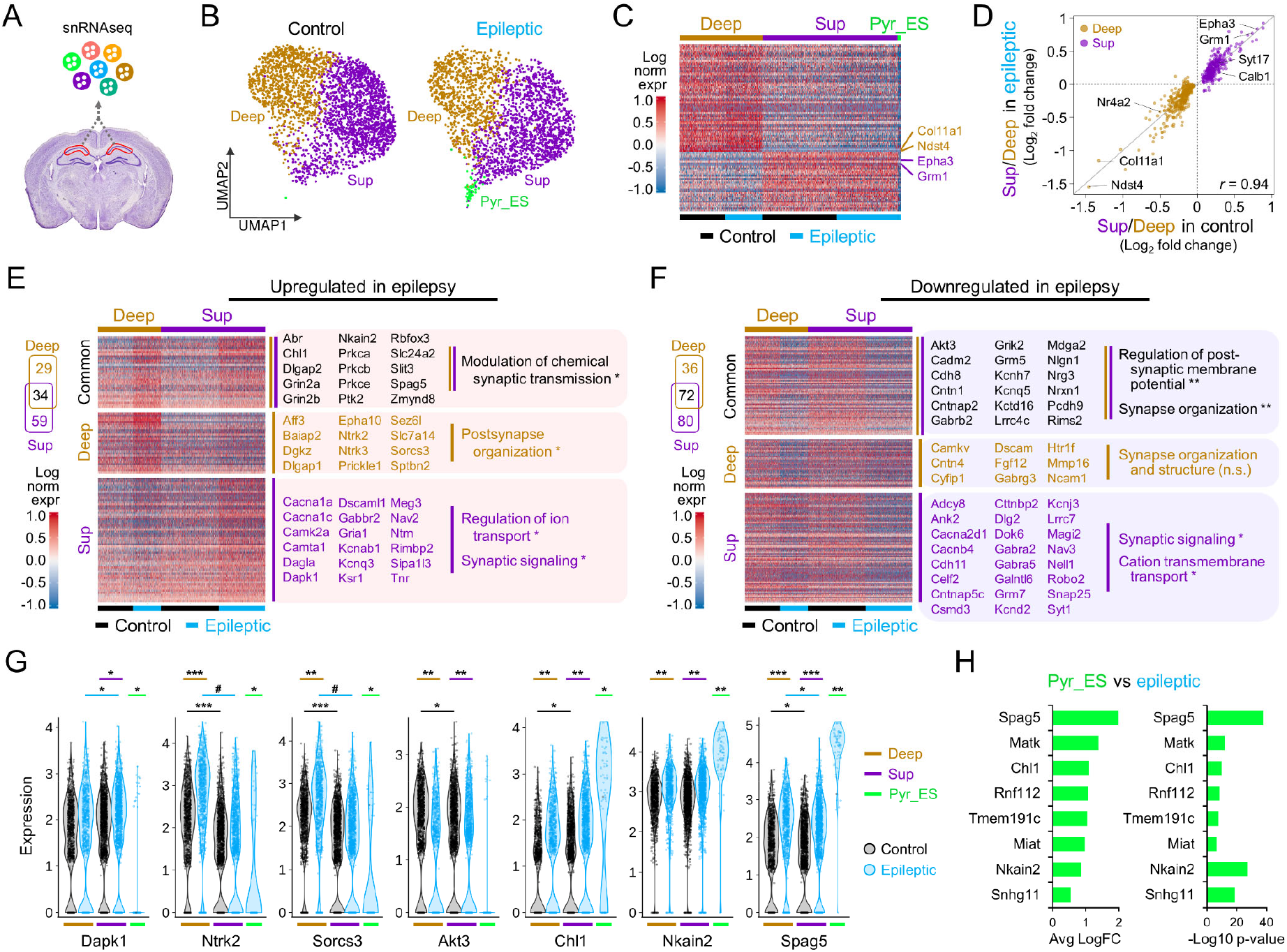
Single nucleus RNAseq profiling of the normal and epileptic CA1 area. **A**, Experimental schema of intact nuclei isolation and massive parallel droplet encapsulation for single-nucleus transcriptional profiling. Nuclei were isolated from CA1 region of adult mice and purified by flow cytometry for single nucleus RNAseq (snRNAseq). **B**, UMAP plots of CA1 pyramidal neuronal subtypes segregated by condition (control and epileptic). Pyr_ES: epilepsy-specific. **C**, Heatmap showing normalized expression (normalized log transformed UMIs) for principal gene markers for deep (Deep), superficial (Sup) neurons in control and epileptic mice (96 enriched genes: 62 deep, 34 sup; absolute LFC>0.25; min. pct>0.5; Adj p-value<1e-30). Note the presence of a subset of cells with differential gene expression corresponding to the epilepsy-specific cell population (Pyr_ES, green). **D**, Scatter plot showing significantly enriched genes in superficial and deep CA1 neurons in control mice and their relative level of enrichment in epileptic animals (Adj p-value<0.05; 493 genes). Gene expression levels of CA1 neuronal subtype-specific enriched genes are preserved in epilepsy. **E**, Venn diagram and heatmap of DEGs upregulated by epilepsy (Adj p-value<0.05, Wilcoxon rank sum test). Heatmap shows normalized expression levels (normalized log transformed UMIs) for upregulated DEGs that are common (34 genes) or specific to deep (29 genes) and superficial (59 genes) CA1 neurons. Representative genes and significant GO terms associated to each gene list are highlighted at right. *FDR p<0.05 (Fisher’s exact test). **F**, Same for downregulated genes. **G**, Violin plots showing normalized expression value (normalized log transformed UMIs) of selected genes in deep and superficial neurons in epilepsy (blue) and control (black). Wilcoxon rank sum test *p<0.05; **p<1E-10; ***p<1E-50; #p<1E-100. Expression levels in Pyr_ES cells is also shown (green). **H**, Bar chart of fold change (left) and significance (right) for most upregulated genes in Pyr_ES when compared with epileptic CA1 pyramidal cells (absolute fold change >0.5 and transcript detection in >50% of the nuclei).

**Fig.5.**
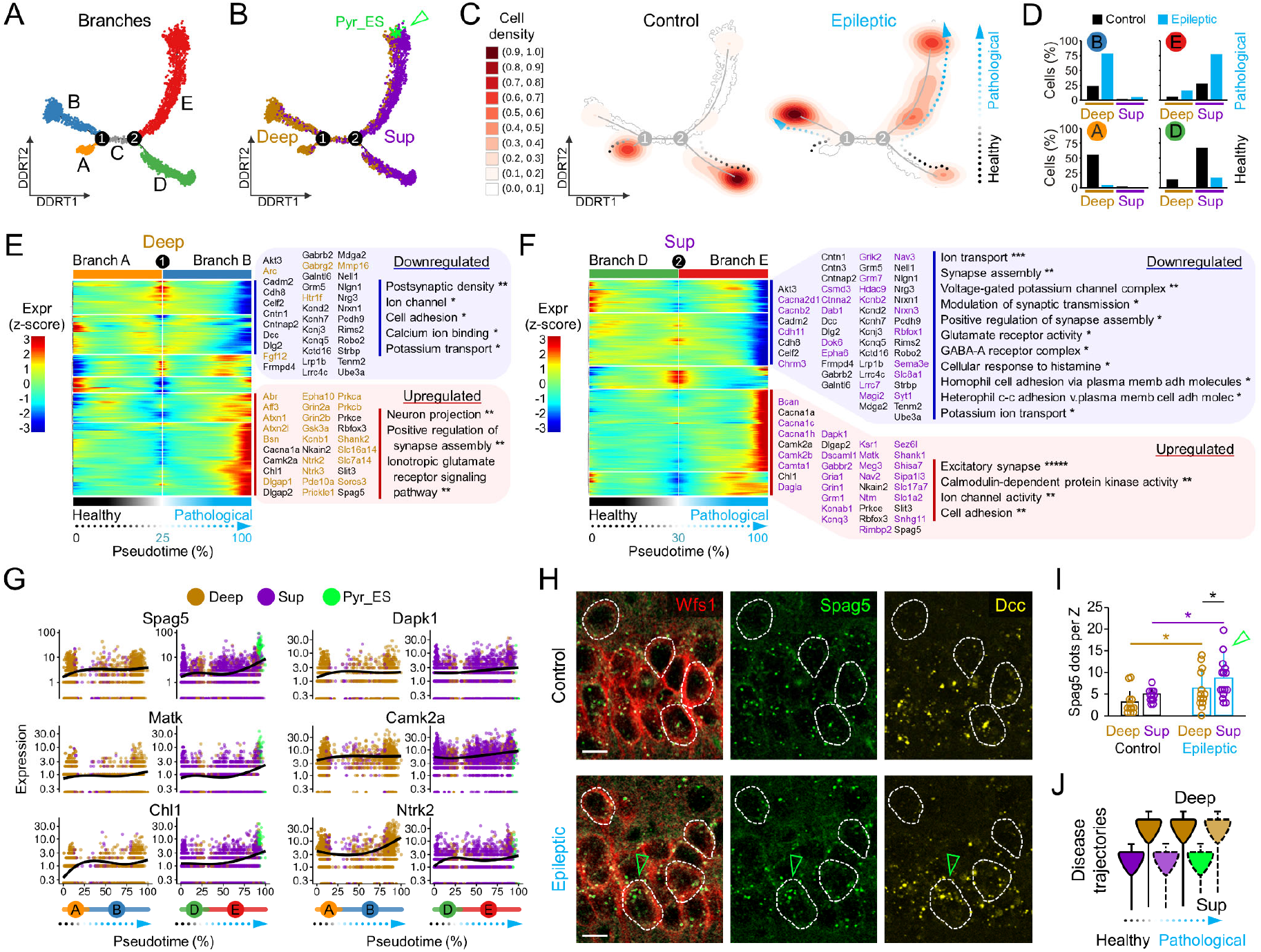
Pseudotemporal ordering of transcriptional changes unfolds separate disease trajectories across CA1 sublayers. **A**, UMAP plot of pseudotemporal trajectory analysis of pyramidal cells subpopulations along DDRTree coordinates uncovers four major branches (A,B,D,E) and segregated from two branching points (1,2). **B**, Same plot as in A showing distribution of pyramidal cells subpopulations (deep, superficial, Pyr_ES) across pseudotime trajectories. Arrowhead points to Pyr_ES cells located at the endpoint of branch E. **C**, Heatmap showing distribution of pyramidal cells subpopulations across pseudotime trajectories by condition (control and epileptic). **D**, Distribution of pyramidal cell subtypes across the trajectory topology. **E**, Heatmap of top-ranked 250 branched expression analysis modeling (BEAM) significant changes through the progression across the trajectory from branch A (basal state) to B (epilepsy) (deep cells). Note deep-specific expression up- (red) and down-regulation (blue) kinetics of the transcripts from the basal healthy state to the epileptic state. **F**, Same for the superficial-specific trajectory from basal state (branch D) to the epileptic condition (branch E). **G**, Examples of gradient progression across disease trajectory for significantly modulated genes in epilepsy related to neuronal survival (*Spag5, Matk, Chl1, Dapk1, Camk2a*, and *Ntrk2*). **H**, Immunostaining against the CA1 pyramidal cell marker Wfs1 and multiplexed RNAscope for *Spag5* and *Dcc* transcripts. Cells with their soma in the confocal plane are outlined. Note significant accumulation of *Spag5* in an epileptic cell (green arrowhead). **I**, Quantification of *Spag5* in deep and superficial CA1 neurons in 3 control and 3 epileptic mice, as counted in one confocal plane per animal (2-way ANOVA effect for group F(1,51)=4.225, p=0.0450 and sublayers F(1,51)=11.521, p=0.0013; *, p<0.05 posthoc t-test). Data from 27 control (14 deep, 14 superficial) and 28 epileptic cells (15 deep, 13 superficial). Green arrowhead indicates cells with high *Spag5* expression. **J**, Schematic summary of disease trajectories reflecting pseudotime progression from control to epilepsy.

We focused on pyramidal CA1 neurons, which represented 80% of cell population (5347 nuclei; 2,934 control and 2,413 epileptic). After two clustering rounds (Suppl.Fig.5E), we identified deep and superficial pyramidal cells from control (1123 deep, 1810 sup) and epileptic samples (878 deep and 1469 sup) and a small population of pyramidal nuclei that were mostly epilepsy-specific (Pyr_ES, 67 nuclei: 66 from epileptic CA1; 1 from control CA1) (Fig.4B). Notably, clustering exhibited consistent distribution of markers for deep (e.g. *Ndst4, Col11a1*) and superficial pyramidal neurons (e.g. *Calb1, Epha3*) (Sup.Fig.5F,G). This specific population of Pyr_ES cells showed total UMI values well above quality threshold (>500) and a number of annotated genes lying within the control and epileptic ranges (Suppl.Fig.5H), excluding potential artifacts. Differential expression analysis of Pyr_ES cells against all detected cells excluding deep and superficial pyramidal neurons, revealed enrichment of pyramidal CA1 marker genes such as *Gria2, Rasgrf1, Camkv* and *Brd9*, along with other transcripts enriched in excitatory neurons including *Hs6st3, Cntnap2, Kcnh7, Kcnip4, and Meg3*, which confirmed their pyramidal nature (Suppl.Fig.5I,J). Similar to our LCM-RNAseq observations at the tissue level, we found consistent correlation of sublayer-specific genes in control and epileptic samples (Fig.4D; Suppl.Fig.6).

First, we sought to identify gene programs underlying the differential vulnerability of deep and superficial CA1 pyramidal cell in response to epilepsy (Adj p-value<0.05). DEGs common to both cell-types were associated with the modulation of synaptic transmission, synapse organization and regulation of membrane potential (Fig.4E,F; common genes in black). Single-nucleus differential expression analysis also revealed striking differences between superficial and deep cells in genes involved in synaptic signaling and ion membrane transport (Fig.4E,F). To exclude effects of different sample size between groups and cell-types, we evaluated the top-ranked 300 DEGs and found similar GO families differentially involved (Suppl.Fig.7A). These include the gamma-aminobutyric acid (GABA) signaling pathway, calcium ion transport, potassium ion transport, regulation of glutamatergic synaptic transmission, axon growth and guidance and synapse assembly. Notably, we also noted cell-type specific changes in genes related to neuronal survival. For instance, we found significant upregulation of the death associated protein kinase 1 (*Dapk1*) specifically in superficial CA1 neurons (*36*) (Fig.4G). Conversely, genes associated to pro-survival receptors TrkB (*Ntrk2*), TrkC (*Ntrk3*), and the sortilin-related VPS10 domain containing receptor 3 (*Sorcs3*) were exclusively upregulated in deep cells (*37, 38*) (Fig.4G). Other genes regulating neuronal vulnerability to insults were found deregulated in both populations of CA1 pyramidal neurons. These include the anti-apoptotic genes *Akt3, Chl1* and *Spag5*, and the pro-apoptotic gene *Nkain2* (*39–41*) (Fig.4G).

We next focused on the small cluster of epilepsy-specific Pyr_ES cells (Fig.4B, green). When compared against control pyramidal cells, Pyr_ES cells showed transcriptional dysregulation of the GO families that were altered in superficial and deep CA1 neurons in epilepsy (Suppl.Fig.7B). To avoid confounding effects caused by the small sample size, we focused in those transcripts identified as differentially expressed between Pyr_ES and all other epileptic cells, that were well represented in both populations (i.e. detected in at least 50% cells per group; min. pct1>0.5 and min. pct2>0.5). Notably, many of these transcripts were upregulated more largely in Pyr_ES neurons as compared to other epileptic cells, including the previously mentioned apoptotic-related genes *Spag5, Chl1* and *NKain2* (Fig.4H; Suppl.Fig.7B,C).

Therefore, our analysis confirmed profound transcriptional changes related to neuronal excitability and neurodegeneration in epileptic CA1 pyramidal neurons. Notably, these functions appear to be segregated across deep and superficial cell-types consistent with electrophysiological, LCM-RNAseq and histological data. Notably, snRNAseq analysis revealed disease-associated genes that were highly cell-type specific, including some with key functions in neuronal survival. Hence, epileptic-induced cell-type-specific regulation of genes with key roles in synaptic plasticity and survival is a potential molecular link between differential activity burden and vulnerability across CA1 radial axis leading to regionalized hippocampal sclerosis.

### Pseudotemporal ordering reveals sublayer-specific neurodegenerative progression

Based on data above we speculated that epilepsy-related responses and degenerative signals might evolve distinctly across cell-types and sublayers. This may be especially critical for cell death programs accompanying hippocampal sclerosis. To glean insights into these transitional states, we used manifold learning leveraged in nearest-neighbor information to automatically organize cells in trajectories along a principal tree reflecting progression of associated biological processes (*42*).

Low dimensional embedding of the automatically learnt underlying trajectory produced a spanning tree revealing a topological structure with four main branches (A, B, D, E) and two bifurcation points (1, 2; Fig.5A). Most nuclei at branches A and B were from cells identified as deep, whereas branches D and E were populated mostly by superficial cells (Fig.5B). Interestingly, control and epileptic cells distributed differently. Branches A and D were mostly populated by control pyramidal cells, whereas branches B and E contained the vast majority of epileptic cells (Fig.5C,D).

The trajectory topology suggested that the transcriptional state of single cells progress along sublayer-specific disease trajectories from a basal state (branches A, D) to the epileptic condition (branches B, E) (Fig.5C). Notably, epilepsy-specific Pyr_ES cells were retrieved at the end of the superficial epileptic E path, suggesting that epileptic Pyr_ES cells may be superficial neurons at a terminal pathological state (Fig.5B; green arrowhead).

To identify mechanisms underlying single-cell transcriptional changes from health to disease we performed quantitative comparison of gene expression kinetics across trajectories in each cellular population (DDRTree method) (*43*). By comparing the top-ranked 250 DEGs in deep (max FDR = 8.94E-4; likelihood ratio test) versus superficial cells (max FDR=5.8E-19), we found about 50% of genes (124 genes) associated to epilepsy carrying significant cell-type specific trajectories. Consistent with our previous results, we found that genes deregulated in epilepsy in superficial CA1 neurons show tight functional association among them, leading to high proportion of significantly enriched ontology terms as compared to genes deregulated in deep CA1 neurons (Fig.5E,F). Again, significantly enriched ontology terms were related to structural and functional neuronal plasticity. Interestingly, pseudotemporal kinetics also revealed marked differences in the progression across sublayers of genes with important functions in regulating neuronal survival. For instance, gene transcript levels of the death associated protein kinase 1 (*Dapk1*) showed robust increase in superficial epileptic cells, whereas gene expression levels of the pro-survival receptors TrkB (*Ntrk2*) and TrkB (*Ntrk3*) along with other genes playing key functions in neurons such as *Sorcs3* and *Aff3* were upregulated specifically in deep pyramidal cells (Fig5G, Sup.Fig.7D; Sup.Table3). Other genes also related to neuronal vulnerability such as *Camk2a* and *Nrg3* displayed similar changes along disease trajectory in both cell types. Strikingly, the subset of apoptotic-related genes *Spag5, Matk* and *Chl1* reached their maximal expression level in Pyr_ES at the end of the transcriptional trajectory in superficial cells (Fig.5G, green).

To confirm these trends *in situ* we combined the single-molecule amplification method RNAscope with immunostaining against the CA1 pyramidal cell marker Wfs1 in coronal brain sections of control and epileptic mice (Fig.5H; Sup.Fig.7E). For RNAscope we selected *Spag5*, which exhibited the largest changes in epilepsy (Fig.5G; see Fig.4H), and the superficial gene marker *Dcc* which is not affected by epilepsy (Supp.Fig.6A; Supp.Fig.7F-H). We found different distribution of *Spag5* dots in sections from control and epileptic mice with significant effects for groups (2-way ANOVA F(1,51)=4.225, p=0.0450) and sublayers (F(1,51)=11.521, p=0.0013; no interaction, all cells counted in one confocal section per animal; 3 control, 3 epileptic) (Fig.5I). Consistent with trajectory analysis, a small subset of pyramidal cells exhibited large levels of Spag5 dots roughly resembling expression distribution from snRNAseq data (Fig.5H, green arrowhead; see distribution in Fig.4G), giving support to the idea that these cells represent a later stage in neurodegeneration (Fig.5J).

## Discussion

Our work identifies sublayer-specific transcriptional changes in experimental hippocampal sclerosis. Using a combination of techniques, we found that acquired TLE involves heterogeneous biological processes running across deep and superficial CA1 pyramidal cells. Our data suggest that epileptogenesis and the accompanying hippocampal sclerosis are evolving processes that affect the transcriptional state of neurons in a cell-type specific manner.

Hippocampal sclerosis is a heterogeneous histopathological entity (*2*). The prognosis value of certain subtypes has been largely debated but emerging data suggest it deserves further consideration in light of cell-type specificity (*12*). Superficial and deep CA1 pyramidal cells are differently innervated by local circuit GABAergic interneurons, express different neuromodulatory receptors, project differentially to cortical and subcortical regions and participate distinctly of hippocampal oscillations (*17, 19, 20*). The more sustained excitability of superficial pyramidal cells (in terms of post-event depolarization and increased firing rate) and their consistent stronger expression of the immediate-early gene associated protein c-Fos after individual seizures, suggest they undergo larger activity burden as compared with deep cells (Fig.1). While our snRNAseq analysis identified many transcripts differentially regulated between deep and superficial CA1 neurons that may underlie pro-epileptogenic changes in intrinsic excitability, microcircuit mechanisms are key contributors to these differences. First, perisomatic inhibition by parvalbumin (PV) basket cells is remarkably higher in deep than superficial cells (*20, 44*). Second, superficial cells are more likely to be driven by presynaptic CA3 activity than deep cells (*44*). Similarly, direct inputs from layer III pyramidal neurons in the medial entorhinal cortex are biased by deep CA1 cells, whereas projections from the lateral entorhinal cortex are stronger in superficial (*45*). Given loss of medial layer III inputs and sprouting of lateral inputs together with specific cannabinoid type-1 receptors pathways (*46, 47*), it is very much likely all these changes contribute to more sustained activation of superficial cells.

Transcriptional dysregulation is a central feature of most neurodegenerative diseases. Our transcriptional profiling in laser-microdisected CA1 deep and superficial sublayers (LCM-RNAseq) revealed strong regionalized responses to acquired epilepsy. We found significantly larger transcriptional changes of GO terms associated with the immune and inflammatory processes, cytokine production and programmed cell death in the superficial sublayer (Fig.2). This includes *Cx3cr1, P2ry12* and *Il18* amongst others previously described genes to play major roles in microglial activation and neurodegeneration associated to epilepsy (*48-50*). For deep pyramidal sublayers transcriptional changes were milder in general and mostly involved deregulation of genes associated with changes of excitability, homeostatic regulation and synaptic signalling, such as *Kcnq2, Kcnq5, Cntnap2*, and *Gabrb2* (*51-53*). Many of these changes were confirmed with snRNAseq analysis (Fig.4), which disclosed significant upregulation of the death associated protein kinase 1 (*Dapk1*) specifically in superficial CA1 neurons (*36*) while genes associated to pro-survival receptors TrkB (*Ntrk2*), TrkC (*Ntrk3*), and the sortilin-related VPS10 domain containing receptor 3 (*Sorcs3*) were mostly upregulated in deep cells (*37, 38*).

By leveraging on unsupervised branched expression learning, we identified deviating transcriptional disease trajectories in deep and superficial pyramidal cells (Fig.5). Some of the GO terms involved suggest that transition from the normal to the epileptic condition in deep cells is more likely associated to changes of synaptic signalling and excitability (i.e. they reflect microcircuit alterations), whereas superficial cells rather move differentially along transcriptional changes impacting on cell-cell communication (i.e. neuron-microglia interactions). Such different latent trends might actually suggest that deep and superficial cells are at different stages in the neurodegenerative process or being affected by pathways leading to differential vulnerability. For instance, while we found deregulated expression of genes associated with apoptotic pathways that were common to both cell types in response to epilepsy (i.e. *Akt3, Nrg3 and Camk2a*) (*39, 41, 54–56*), genes related to cell death were found deregulated specifically in superficial or deep neurons. In superficial cells, epilepsy resulted in decreased expression of brain derived neurotrophic factor (*Bdnf*), and increased transcript levels of pro-apoptotic pathway activator Death-associated protein kinase 1 (*Dapk1*) (*36*). Conversely, in deep pyramidal neurons, epilepsy led to robust increase in the expression of the gene encoding the pro-survival receptors TrkB (*Ntrk2*) and TrkC (*Ntrk3*)(*37, 38*). Interestingly, pseudotime analysis mapped fewer control cells along the epileptic branches in both sublayers suggesting that latent sublayer specific transcriptional processes might actually be running along life (*57*).

Our snRNAseq analysis also disclosed an epilepsy-specific pyramidal cell population, Pyr_ES, which accumulated at the end of the superficial trajectory branch (Fig.5B, arrowhead). This subset of pyramidal cells displayed remarkable expression of neurodegeneration related transcripts such as *Cdk5, Ckb, Matk, Chl1, and Spag5* (*58*). The presence of cells with very high number of *Spag5* molecules in the superficial sublayer of the epileptic CA1 was confirmed by combined immunofluorescence and single-molecule amplification methods. Thus, our results indicate that genes more largely dysregulated in epilepsy are specifically expressed by vulnerable subset of pyramidal neurons undergoing later stages of neurodegeneration by the time of sampling. We propose that Pyr_ES cells reflect the accumulated pro-epileptic transitional changes leading to epileptogenesis and neurodegeneration, as suggested by their extreme location along disease trajectories.

Altogether, our results identify previously unobserved heterogeneity in the neuronal patterns of activity of deep and superficial CA1 pyramidal neurons in epilepsy, discover specific gene expression signatures across CA1 deep-superficial sublayers that are associated to neuronal loss and hippocampal sclerosis, reveal disease trajectories of deep and superficial CA1 pyramidal neurons in epilepsy, and uncover the underlying transcriptional programs. By dissecting the transcriptional landscape across CA1 sublayers in epilepsy, our work offers new insights into the mechanisms regulating epileptogenesis and highlights the importance of leveraging on cell type specificity to better understand the phenotypic complexities accompanying hippocampal sclerosis in epilepsy.

## Materials and Methods

All experimental protocols and procedures were performed according to the Spanish legislation (R.D. 1201/2005 and L.32/2007), the European Communities Council Directives of 1986 (86/609/EEC) and 2003 (2003/65/CE) for animal research, and were approved by the Ethics Committee of the Instituto Cajal.

### Epilepsy model

Adult male Wistar rats (180–200 g), as well as wild-type C57 and transgenic mice (20-25 g), were treated with multiple intraperitoneal injections of kainate (5 mg/kg) at hourly intervals until they reached *status* epilepticus. Transgenic mouse lines included the Thy1.2-G-CaMP7-DsRed2 (c57BL/6J-Tg(Thy1-G-CaMP7-DsRed2)492Bsi, stock RBRCO6579, RIKEN), the Calb1-Cre (Jackson Lab, stock No:023531; Calb1-2A-dgCre-D) and the Calb1-Cre line crossed with the tdTomato reporter line (Jackson Lab, stock No:007905; Ai9). The *status* was defined as a condition of continuous seizures lasting longer than 30 min. In a subset of rats, the lithium-pilocarpine model was used. These rats were i.p. injected with pilocarpine hydrochloride 12–24 hr after the injection of lithium chloride (127 mg/kg, i.p.). Between one and four doses of 10 mg/kg pilocarpine were injected every 30 min until the *status* epilepticus was reached. Diazepam (4 mg/kg) was injected 1 h post-*status* to stop convulsions in all animals. They received intraperitoneal injections of 5% dextrose in saline (2.5 ml) and their diet was supplemented with fruit and powder milk during the following 2–3 days. After 3 days, animals behaved normally and were housed individually. Control animals were injected with saline and received treatments similar to epileptic animals.

All experiments started after 6 weeks post-*status* and extended up to 25 weeks post-*status*, well into the chronic epileptic phase. Seizures were observed in all animals during handling or recorded electrophysiologically in a subset of animals. In some epileptic animals, we aimed to induce convulsive seizures using high-pitched sounds (random pulses of 95-100 dB and 1-20 sec duration at 0.05-1 Hz during 10 min). A convulsive motor seizure was defined according to clinical criteria, as sustained and repeated forelimbs automatisms with or without falling. Control animals did never exhibit convulsive seizures using this protocol.

### EEG recordings and analysis

To evaluate epileptogenesis, some rats and mice were implanted with either intracranial 16-channel silicon probes or skull EEG-grid of 32-channels (Neuronexus) under isoflurane anesthesia (1.5–2% mixed in oxygen 400–800 ml/min). Jeweler’s screws were inserted into the skull for providing additional anchoring and reference/ground connections (over the cerebellum). The implant was secured with dental cement. Animals were recovered from anesthesia and returned to home cages.

For recordings, EEG signals were pre-amplified (4x gain) and recorded with a 32-channel AC amplifier (Axona), further amplified by 100, filtered by analog means at 1Hz to 5 kHz, and sampled at 20 kHz/channel with 12 bits precision. EEG recordings were synchronized with a ceiling video camera (30 frames/sec) to track the animal position in space.

Analysis of electrophysiological signals was implemented in MATLAB 9.3 (MathWorks). EEG signals from the frontal and parietal electrodes were used to identify theta periods during running (band-pass 4-12 Hz) and periods of immobility characterized by low frequency delta activity (0.5-4Hz). Forward-backward-zero-phase finite impulse response (FIR) filters of order 512 were used to preserve temporal relationships between channels and signals. Spectral values fitted to 1/f were similar between groups for frequencies >150 Hz. HFO events were defined from the bandpass filtered signal (80-120 Hz) by thresholding (>4 SDs). The power spectra were evaluated in a window of ± 0.2 ms around each detected event. Time-frequency analysis was performed by applying the multitaper spectral estimation in sliding windows with 97.7% overlap and frequency resolution of 10 Hz in the 90-600 Hz frequency range. HFO activity was evaluated as the power integral in the 80-120 Hz band.

We used a combination of features to identify HFO events automatically and to classify them in different categories (*23*) (Supp.Fig.1). First we identified large amplitude transient (<100 ms) discharges using LFP and current-source-density signals (CSD, i.e. the second spatial derivative) at the SR and SLM. Second, we identified HFO events at the SP by frequency thresholding over 100 Hz. Then we used spectral indices such as entropy and fast ripple indices from candidate HFO events at SP together with amplitude information from LFP and CSD signals at SR to classify events as: a) SPW-ripples (low amplitude; 100-150 Hz); b) SPW-fast ripples (medium amplitude, >150 Hz) and IID-HFO (larger amplitude, >100 Hz). Events not meeting criteria were left unclassified. Surface EEG recordings from mice were analyzed similarly, by using channels over the dorsal hippocampus or at the frontal cortex to identify HFO events and channels over the parietal cortex using selected segments of the EEG.

### In vivo recording and labeling of single cells

Rats were anesthetized with urethane (1.2 g/kg, i.p.), fastened to the stereotaxic frame and warmed to keep their body temperature at 37°. Two bilateral craniotomies of ~1 mm diameter were performed for CA3 stimulation (AP: −1.2 mm, ML: 2.9 mm) and CA1 recordings (AP: −3.7 mm; ML: 3 mm). The dura was gently removed, the *cisterna magna* was drained and the craniotomy covered with warm agar to reinforce stability.

A 16-channel silicon probes (NeuroNexus Tech; 100 μm interspaced, 413 μm^2^ contact) was advanced perpendicular along the CA1-DG-CA3c axis guided by extracellular stimulation and electrophysiological hallmarks. Extracellular signals were pre-amplified (4x gain) and recorded with a 16(32)-channel AC amplifier (Multichannel Systems), further amplified by 100, analogically filtered at 1Hz to 5 kHz, and sampled at 20 kHz/channel with 12 bits precision with a Digidata 1440. Concentric bipolar electrodes were advanced 3.5 mm with 30° in the coronal plane to stimulate CA3. Stimulation consisted of biphasic square pulses (0.2 ms duration, 0.05-1.2 mA every 5 s). A subcutaneous Ag/AgCl wire in the neck served as reference. Recording and stimulus position was confirmed by post hoc histological analysis.

Intracellular recording and labelling were obtained in current-clamp mode using sharp pipettes (1.5 mm/0.86 mm outer/inner diameter borosilicate glass; A-M Systems, Inc) were filled with 1.5 M potassium acetate and 2% Neurobiotin (Vector Labs, Inc; 50-100 MΩ) (*59*). Signals were acquired with an intracellular amplifier (Axoclamp 900A) at 100x gain. The resting potential, input resistance and amplitude of action potentials was monitored all over the course of experiments.

After data collection, Neurobiotin was ejected using 500 ms depolarizing pulses at 1-3 nA at 1 Hz for 10-45 min. Animals were perfused with 4% paraformaldehyde (PFA) and 15% saturated picric acid in 0.1 M, pH 7.4 phosphate buffered saline (PBS). Brains were postfixed overnight at room temperature (RT), washed in PBS and serially cut in 70 μm coronal sections (Leica VT 1000S vibratome). Sections containing the stimulus and probe tracks were identified with a stereomicroscope (S8APO, Leica). Sections containing Neurobiotin-labeled cells were localized by incubating them in 1:400 Alexa Fluor 488-conjugated streptavidin (Jackson ImmunoResearch 016-540-084) with 0.5% Triton X-100 in PBS (PBS-Tx) for 2 hours at room temperature (RT). To evaluate morphological features of single cells, sections containing the somata of recorded cells were processed with Triton 0.5% in PBS, blocked with 10% fetal bovine serum (FBS) in PBS-Tx and incubated overnight at RT with the primary antibody solution containing rabbit anti-Calbindin (1:500, CB D-28k, Swant CB-38) or mouse anti-Calbindin (1:1000, CB D-28k, Swant 300) antibodies with 1% FBS in PBS-Tx. After three washes in PBS-Tx, sections were incubated for 2 hours at RT with appropriate secondary antibodies: goat anti-rabbit Alexa Fluor 633 (1:500, Invitrogen, A21070), and goat anti-mouse Alexa Fluor488 (Jackson Immunoresearch, 115-545-003) or goat anti-mouse Rhodamine Red (1:200, Jackson ImmunoResearch, 115-295-003) in PBS-Tx-1% FBS. Following 10 min incubation with bisbenzimide H33258 (1:10000 in PBS, Sigma, B2883) for nuclei labelling, sections were washed and mounted on glass slides in Mowiol (17% polyvinyl alcohol 4-88, 33% glycerin and 2% thimerosal in PBS).

All morphological analyses were performed blindly to electrophysiological data. The distance from the cell soma to radiatum was measured from confocal images using information from Calbindin and bisbenzimide staining. All pyramidal cells included in this study were localized within the CA1 region. Calbindin immunostaining was used to estimate the width of the superficial sub-layer from the border to the stratum radiatum. Superficial cells were defined based on the location of the soma within the calbindin sublayer, independently on their immunoreactivity (*60*). The border with radiatum was estimated for each section and the distance from the recorded cell somata was measured using ImageJ (NIH Image).

### Analysis of intracellular single-cell recordings

Analysis of electrophysiological data was performed using routines written in MATLAB 7.10 (MathWorks). Local field potential (LFP) recorded from sites at the stratum radiatum and lacunosum moleculare were low-pass filtered at 100 Hz to identify sharp-waves and interictal discharges using forward-backward-zero-phase finite impulse response (FIR) filters of order 512 to preserve temporal relationships between channels and signals. LFP signals from sites at the stratum pyramidale were bandpass filtered between 100-600 Hz to study HFOs. For sharp-waves and interictal discharges, filtered signals were smoothed by a Gaussian kernel and events were detected by thresholding (>3 SDs). Interictal discharges were separated from sharp-waves by amplitude and correlation across channels. For HFO detection, the bandpass-filtered signal was subsequently smoothed using a Savitzky-Golay (polynomial) filter and events detected by thresholding (>2 SDs) after artifact and noise rejection. All pairs of detected events were visually confirmed and aligned by the peak of the accompanying sharp-wave and/or interictal discharge. Time-frequency analysis of HFO events was performed by applying the multitaper spectral estimation in sliding windows with 97.7% overlap and frequency resolution of 10 Hz in the 90-600 Hz frequency range (only the 100-600 Hz range is shown) to data sweeps aligned by sharp-wave ripple events (± 1sec). Membrane potential responses of single-cells were evaluated in peri-event plots before (−200 to −150 ms), during (±50 ms) and after (150 to 200 ms) HFO events.

Passive electrophysiological properties (input resistance, membrane decay and capacitance) of neurons recorded intracellularly in vivo were measured using 500 ms currents step in current-clamp mode. Cells with intracellular action potential amplitude smaller than 40 mV were excluded. Resting membrane potential and input resistance were estimated by linear regression between baseline potential data and the associated holding current. Intrinsic firing properties, including action potential threshold, half-width duration and AHP were estimated from the first spike in response to depolarizing current pulses of 0.2 nA amplitude and 500 ms duration. The sag and maximal firing rate was calculated from current pulses of ±0.3 nA amplitude. A bursting index was defined as the ratio of the number of complex spikes (minimum of 3 spikes <8 ms inter-spike interval) over the total number of spikes recorded during theta activity.

### cFos immunostaining and analysis

To evaluate immediate-early gene expression associated to sound-induced convulsive seizures, animals were perfused 1 hour after and their brains cut in 70 μm coronal sections. Selected sections were stained against c-Fos using a polyclonal antibody at 1:250 (Santa Cruz Biotechnology sC-52) and bisbenzimide. Using one 20x confocal moisaic per animal, we quantified cFos intensity at CA1 pyramidal cells by delineating single-cell nuclei stained with bisbenzimide in one confocal plane (ImageJ). The mean intensity of cFos signal from each cell was then normalized by subtracting the background (set at 0) and dividing by the maximal positive signal in the mosaic, which was always at granule cells (set at 1). No significant differences of background were observed across sections. Delineated cells were ranked by their distance to radiatum to classify them as deep or superficial, according to standard measurements of Calb1-layer thickness.

### Laser capture microdissection (LCM) and RNA isolation

Brains from 3 control and 3 epileptic rats were dissected, longitudinally cut in half (separating both hemispheres), wrapped in aluminum foil and immediately frozen in liquid nitrogen. To avoid circadian effects on gene expression all samples were collected in the morning before noon and conserved at −80°C until use. The hippocampal region of each hemisphere was cut in 20μm slices in a cryostat (Leica) (chamber temperature: −20°C; block temperature: −30°C) and placed on 1.0mm PEN-membrane covered slides (Carl Zeiss). Slides were conserved at −20°C until use. Right before microdissection, slides were dried with vapor of liquid nitrogen. The CA1 cell layer was microdissected with a Leica 6000 laser microdissector through a 40x non-oil immersion objective to obtain cell bodies of superficial and deep sublayers separately (Fig.2A). Microdissected deep and superficial areas were collected in different empty caps of 0.5ml Eppendorf tubes. After microdissection, samples were processed following ARCTURUS PicoPure RNA Isolation Kit (Thermo Fisher Scientific) instructions in order to extract and isolate total RNA. Briefly, 50μl of extraction buffer was added into the cap, incubated at 42°C for 30min, centrifuged at 800×G for 2min and stored at −80°C. The same volume of 70% ethanol was added to the cell extract and the mixture was pipetted into a pre-conditioned RNA purification column. The column was centrifuged 2min at 100×G and 30sec at 16000×G, and washed with 100μl of Wash Buffer 1. To completely eliminate DNA, the purification column was treated with 40μl of DNAse (diluted 1/8 in RDD Buffer) (Qiagen), incubated 15min and centrifuged at 8000×G for 15sec. Then the column was washed twice with 100μl of Wash Buffer 2, and centrifuged at 8000×G after the first wash and at 16000×G after the second one. Finally, RNA was eluted into a new 0.5μl Eppendorf tube by adding 11μl of elution buffer onto the column membrane, incubating the column for 1min at room temperature, and centrifuging the column for 1min at 1000×G and at 16000×G immediately after. Total RNA samples were stored at −80°C. RNA integrity number (RIN) was similar in control (4.7 ± 0.8) and epileptic rats (5.7 ± 0.5; P=0.07; 3 replicates x 2 sublayers per group), as well as for deep (5.2 ± 0.9) and superficial sublayers (5.3 ± 0.1; P=0.81; n=6 replicates per sublayers).

### LCM-RNAseq library construction and sequencing

RNA preparation for sequencing deep and superficial CA1 sublayers from control and epileptic rats was performed as described in (*61*). The twelve samples were sequenced according to manufacturer instructions in a HiSeq2500 sequencer (Illumina, Inc). Libraries were strand specific (reverse) and sequenciation was performed in paired-end configuration with a read length of 75bp. Library size of read pairs for the different samples analyzed was between 47 and 59 Million reads. RNAseq data can be accessed at the GEO repository (GSE143555; reviewer token: ehitaysejlwdjad).

### LCM-RNAseq data analysis

Alignment quality control of sequenced samples (LCM-RNAseq) was assessed with FastQC (v.0.11.3) (Babraham Institute) and RNAseq tracks were visualized using IGV (v.2.3.57) (*62*). LCM-RNAseq reads were mapped to the rat genome (Rnor_6.0.83) using STAR (v.2.5.0c) (*63*), and files were further processed with Samtools (v.0.1.19). Aligned reads were counted to gene transcripts using HTSeq (v.0.6.1) (*64*). Differential expression analysis was performed using DESeq2 (v.1.10.0) (*65*) of the bioconductor suite (*66*) in the R (v.3.2.2) statistical computing platform. The experimental design consisted in two factors (treatment and anatomical area) and there was also grouped samples (samples from different anatomical areas (minus (deep) and plus (superficial) that were obtained from individual mice). Genes were considered differentially expressed at Benjamini-Hochberg (BH) Adj p-value<0.05 and absolute log2 fold change>0.3 (*67*), except otherwise specified. GO analysis was performed using DAVID (v.6.8) bioinformatics platform (*68*).

### LCM-RNAseq single-cell informed data analysis

Single-cell RNAseq data from (*14*) was reanalyzed with consensus cluster SC3 algorithm (*69*). From the original 3005 cells, pyramidal and interneurons were removed from somato-sensory cortex, remaining 2442 cells. Remaining cells were reanalyzed downstream with SC3. Clustering stability was optimal for 6 clusters. After that, one cluster presented high heterogeneity (mixed population cluster), and was reanalyzed with the optimal clustering stability (5 clusters). Marker genes were tested for every cluster with Wilcoxon signed rank test. Top 10 genes with the area under the receiver operating characteristic (ROC) curve power >0.85 and with the Adj p-value<0.01 from both cluster analyses were selected, with a total of 69 bona-fide population markers. Based on the markers, populations were fused/splitted and 3 populations were isolated in the first clustering (pyramidal neurons, interneurons and oligodendrocytes) and 4 more in the second (astrocytes, endothelial cells, microglia and mural cells). Sixteen outlier cells were removed by total_counts, total_features or pct_counts_spike criteria, remaining 2,426 cells with a high correspondence with the original classification: Astrocytes (155), Endothelial (177), Interneurons (174), Microglia (85), Mural (56), Oligodendrocytes (804) and Pyramidal (975). To perform an informed analysis leveraged on the subsetted and reanalized data from (*14*), the LCM-RNAseq was analyzed using the 69 markers to identify genes that correlated with cell type. Furthermore, the relevant dimensions (significant or top genes) from LCM-RNAseq data were used over the subsetted and the reanalized dataset, to isolate the contribution of different cell-types across the control and epileptic conditions.

### Single-nuclei isolation

We accurately isolated single nuclei from 2 control and 2 epileptic Calb1::CrexTdtomato young adult mice. We used these animals to facilitate identification of the CA1 region under a fluorescent scope. Animals were sacrificed 12 weeks after the kainate/saline administration by cervical dislocation and brains were dissected and cut in 300um thick slices in a vibratome covered by ice cold HBBS1x. As in LCM studies, samples were collected before noon to avoid circadian effects. The dorsal CA1 region were manually dissected from 4 consecutive slices and put altogether into 400uL of ice-cold MACS buffer (0,5%BSA, 2mM EDTA, PBS1x). CA1 portions were transferred to a dounce homogeneizer (20404 Lab Unlimited) containing 400uL of MACS buffer and were homogeneized 12-15 times each with the pestle. The cell suspension was transferred to a 2mL Eppendorf tube and centrifuge 15min 500G 4°C. Cell pellets were resuspended in 500uL of lysis buffer (10mM Tris-HCL, 10mM NaCl, 3mM MgCl2, 0,1% IGEPAL) and kept 5 minutes on ice. Samples were then spun down at 500G for 30 min in a pre-chilled centrifuge. The nuclei pellet was resuspended in PBS1x 1%BSA and then 15,000 nuclei were purified by flow cytometry in a BD FACS Aria III. The whole process was carried out at 4°.

### Single-nucleus RNA sequencing

Purified intact nuclei from mouse hippocampal CA1 area were processed through all steps to generate stable cDNA libraries. For every sample, 15,000 nuclei were loaded into a Chromium Single Cell A Chip (10x Genomics) and processed following the manufacturer’s instructions. Single-nuclei RNAseq libraries were prepared using the Chromium Single Cell 3 ‘ Library & Gel Bead kit v2 and i7 Multiplex kit (10x Genomics). Pooled libraries were then loaded on a HiSeq2500 instrument (Illumina) and sequenced to obtain 75 bp paired-end reads following manufacturer instructions. On sequencing depth, 262 million fragments were generated for the control condition and 296 for the epileptic dataset. Libraries reached a sequencing saturation of 86.9% for control and 91.2% for epilepsy condition. snRNAseq data can be accessed at the GEO repository (GSE143560; reviewer token: gtkhqgegnjgldur).

### Single-nucleus RNA sequence analysis

Quality control of sequenced reads was performed using FastQC (Babraham Institute). Sequenced samples were processed using the Cell Ranger (v.2.2.0) pipeline (10x Genomics) and aligned to the CRGm38 (mm10) mouse reference genome customized to count reads in introns (pre-mRNA) over the Ensembl gene annotation version 94. Barcodes with total unique molecular identifier (UMI) count >10% of the 99^th^ percentile of the expected recovered cells were selected for further analysis. Using this criterion, we retrieved 3,661 (control), 3,078 (epileptic) high quality nuclei per sample. Mean reads per nucleus were 71,449 (control) and 96,150 (epileptic). Median genes per nucleus were 71,449 (control) and 96,150 (epileptic). Minimum UMI count per nucleus were 710 (control), 540 (epileptic), well above the typical quality standards in single cell/nucleus sequencing. Single-nucleus RNAseq data were subsequently pre-processed and further analysed in R (v.3.4.4) using Seurat (v.2.3.4) (*70, 71*). Filtering parameters were as follows: genes, nCell<5; cells, nGene<200. Data were then normalized using global-scaling normalization (method: LogNormalize, scale.factor=10.000). An initial exploratory analysis was performed on each dataset separately. This analysis retrieved a similar number of clusters in each dataset that were approximately equal in size. We next combine both datasets using the function MergeSeurat. Highly variable genes (HVGs) were detected using FindVariableGenes function with default parameters. Then, normalized counts on HVGs were scaled and centered using ScaleData function with default parameters. Principal component analysis (PCA) was performed over the first ranked 1,000 HVGs, and cluster detection was carried out with Louvain algorithm in FindClusters function, using 20 first PCA dimensions and resolution of 0.6 (the default number in Seurat and the optimal according to cell number, data dispersion and co-expression of previously reported cell markers). Plots of the two principal components of the PCA where cells were colored by dataset of origin excluded the presence of batch effects. This analysis identified 13 clusters. CA1 pyramidal neurons populated three of these clusters: one cluster was enriched in bona fide gene markers of deep cells (*Ndst4, Coll11a1*) whereas a second one was enriched in canonical markers of superficial cells (*Calb1, Epha3*). The third cluster showed a mixed identity. An additional round of clustering segmented this population in three additional clusters that were enriched in deep and superficial markers, respectively, and a third cluster that could not be annotated based on the presence of gene markers of CA1 sublayer neurons. The vast majority of cells within this cluster (66/67) were from epileptic mice (Pyr_ES). Next, the FindMarkers function was used to identify gene markers and to determine the cell populations represented by each cluster. Finally, cell subtypes were manually aggregated based on the presence of canonical markers of known cell types into six distinct major cell types: excitatory neurons (Excit); inhibitory neurons (Inter); oligodendrocytes (ODC); oligodendrocyte progenitor cells (OPCs); microglia (Microglia); astrocytes (Astro). Visualization and embedding were performed using stochastic nearest neighbors (tSNE) (*72*) and uniform manifold approximation and projection (UMAP) (*73*)methods over PCA using the 20 first PCA dimensions. UMAP plots of gene expression show normalized count (UMIs) per nucleus. The equalized expression between fixed percentiles was plotted according to the following criteria: the minimum expression was adjusted to 5% and the maximum expression was adjusted to 95% in all UMAP expression plots. To evaluate effects of epilepsy, datasets from both conditions were merged and HVGs were identified for each dataset as above indicated. Only HVGs that were detected in all datasets were used to perform visualization and embedding as described above. Clustering was performed on merged dataset from both conditions and populations were identified combining these results with clustering information obtained in control and epileptic datasets separately, together with co-expression of population markers. Differential expression analysis (DEA) was used to identify population gene markers. For DEA, the nuclei of each population were contrasted against all the other nuclei in the merged dataset using Wilcoxon Rank Sum test on normalized counts. For epilepsy effect analysis, in the merged dataset, the nuclei of each population from the epileptic dataset were contrasted against all the other nuclei of the same population in control using Wilcoxon Rank Sum test on normalized counts. GO functional enrichment analyses were performed using DAVID (v.6.8) bioinformatics platform (*68*).

### Cell trajectories and pseudotime analysis

The disease pseudotime analysis was performed using Monocle 2 (v.2.8.0) (*42*). First, the Seurat merged dataset was transformed to Monocle object and cells from Pyr_CA1 were subset. The size factor and dispersion of the subset was estimated, and data was normalized and preprocessed. Genes under the minimum level detection threshold of 0.1 and detected in less than 10 cells were filtered with the function setOrderingFilter. Genes defining how a cell progress through a pseudo-time disease trajectory were selected with the function differentialGeneTest (Monocle’s main differential analysis routine). 2579 genes (64,68% of a total of 3987 genes considered as expressed) were significative with FDR<1% for the combination of factors: ~SeuratCluster+Condition, and thus, defined the high dimensional space for pseudotemporal trajectory analysis. Discriminative dimensionality reduction with trees (DDRTree) reduction algorithm learns the principal graph and specifies the trajectory. DDRTree was applied inside the function reduceDimension, and got the default parameters: norm_method = “log”, pseudo_expr = 1, relative_expr = TRUE, auto_param_selection = TRUE (automatically calculate the proper value for the ncenter (number of centroids)) and scaling = TRUE (scale each gene before running trajectory reconstruction). Prior the dimensional reduction, the function reduceDimension also performed a variance-stabilization of the data (because the expressionFamily of the data was negbinomial.size). Finally, the cells were ordered according to pseudo-time with the function orderCells, which added a pseudo-time value and state for each cell; together encode where each cell maps to the trajectory. For enrichment analysis on pseudotime trajectories, top-ranked 250 branched expression analysis modeling (BEAM) significant changes through the progression across the disease trajectory for each sublayer were clusterized and GO enrichment analyses on upregulated and downregulated clusters were performed using DAVID (v.6.8) bioinformatics platform (*68*).

### Cell-type immunostaining and analysis

To evaluate the contribution of different cell-types, control and epileptic rats and mice were perfused with 4% paraformaldehyde (PFA) and 15% saturated picric acid in 0.1 M PBS, pH 7.4. Brains were postfixed overnight and cut in 70 μm coronal sections (Leica VT 1000S vibratome). Sections containing the dorsal-intermediate hippocampus were processed with Triton 0.5% in PBS and blocked with 10% fetal bovine serum (FBS) in PBS-Tx. Sections were incubated overnight at RT with 1% FBS PBS-Tx solution containing primary antibodies against a battery of cell-type specific markers. The list of antibodies include: rabbit anti-calbindin (1:1000, CB D-28k, Swant CB-38) or mouse anti-calbindin (1:500, CB D-28k, Swant 300) to identify superficial CA1 pyramidal cells; rabbit anti-Wfs1 (1:500, Protein Tech, 11558-1-AP) for CA1 pyramidal cells; rabbit anti-Iba1 (1:1000, Wako, 019-19741) for microglia; rabbit anti-GFAP (1:1000, Sigma, G9269) for astrocytes; rabbit anti-Olig2 (1:200, Millipore, AB9610) for oligodendrocytes. After three washes in PBS-Tx, sections were incubated for 2 hours at RT with appropriate secondary antibodies: goat anti-rabbit Alexa Fluor633 (1:200, ThermoFisher, A-21070), and donkey anti-mouse Alexa Fluor488 (1:200, ThermoFisher, A-21202) or goat anti-mouse Rhodamine Red (1:200, Jackson ImmunoResearch, 115-295-003) in PBS-Tx-1%FBS. Following 10 min incubation with bisbenzimide H33258 (1:10000 in PBS, Sigma, B2883) for nuclei labelling, sections were washed and mounted on glass slides in Mowiol (17% polyvinyl alcohol 4-88, 33% glycerin and 2% thimerosal in PBS).

To acquire multichannel fluorescence stacks from recorded cells, a confocal microscope (Leica SP5) with LAS AF software v2.6.0 build 7266 (Leica) was used. For single-cell studies the following channels (fluorophore, laser and excitation wavelength, emission spectral filter) were used: a) bisbenzimide, Diode 405 nm, 415–485 nm; b) Alexa Fluor 488, Argon 488 nm, 499–535 nm; c) Rhodamine Red / Alexa Fluor 568 / Texas Red, DPSS 561nm, 571–620 nm; d) Alexa Fluor 633, HeNe 633 nm, 652–738 nm; and objectives HC PL APO CS 10.0×0.40 DRY UV, HCX PL APO lambda blue 20.0×0.70 IMM UV and HCX PL APO CS 40.0×1.25 OIL UV were used.

### FluoroJade staining

To evaluate neurodegenerating neurons we used coronal sections from epileptic rats perfused at different time points post-*status* (from 2 to 3 weeks). Selected sections were immunostained against Wfs1 followed by FluoroJade staining. To this purpose, sections were pretreated for 5 min with 1% sodium hydroxide in 80% ethanol, followed by 70% ethanol (2 min) and distilled water (2min). Sections were then incubated 10 min in 0.06% potassium permanganate, rinsed in distilled water and immersed into 0.0001% solution of FluoroJade C dye (Sigma AG325) in 0.1% acetic acid (pH 3.5) for 10min. After a brief wash in distilled water, they were mounted on gelatin-coated slides, air-dried, coverslipped with DPX and examined under a confocal microscope as described above. FluoroJade positive cells exhibited bright green fluorescence.

### In situ hybridization analysis

Selected sections from control and epileptic rats were processed for in situ hybridization using standard methods. Briefly, riboprobes were prepared from Rat *Enpp2* cDNA (Image clone ID 7115236) and using RT-PCR from rat adult hippocampus to prepare *Wfs1* (NM_031823.1, from bp 783 to 1631), *Ndst4* (XM_006233274.2, from bp 536 to 1024), *Syt17* (NM_138849.1, from bp 378 to 1118), *Hrt1a* (J05276.1, from bp 730 to 1663) and *Scn7a* (NM_131912.1, from bp 2812 to 3550) cDNAs. Similar riboprobes were used in sections from normal and epileptic mice. Riboprobe hybridization was detected using alkaline phosphatase-coupled anti-digoxigenin Fab fragments (Sigma). Hybridized sections were mounted in glycerol and photographed using a Nikon stereoscope and a DCC Nikon camera.

### RNAscope analysis

Control and epileptic mice were perfused with 4% paraformaldehyde (PFA) and 15% saturated picric acid in 0.1 M PBS, pH 7.4. Brains were post-fixed overnight and cut in 50 μm coronal sections (Leica VT 1000S vibratome). Sections containing the dorsal-intermediate hippocampus were mounted onto SuperFrost Plus microscope slides (10149870, ThermoFisher Scientific). RNAscope Fluorescent Multiplex Assay (Advanced Cell Diagnostics) was carried out according to the manufacturer’s protocols. Briefly, sections were dehydrated at 60°C, follow by ethanol, pretreated with a target retrieval solution (322000, ACD) and protease III (322340, ACD), and co-hybridized with Spag5 (Mm-Spag5, 505691, ACD) and Dcc (Mm-Dcc-C3, 427491, ACD) probes. Finally, the amplification steps (RNAscope Fluorescent Multiplex Detection reagents, 320851, ACD) were followed, using Atto 550 for Spag5 and Atto 647 for Dcc as fluorescent labels. The RNAscope 3-plex positive control probe set (320881, ACD), with probes to Polr2a, PPIB and UBC, was used to confirm preservation of sample RNA. The negative control probe to bacterial DapB (320871) was used to establish non-specific labelling.

Following the RNAscope protocol, sections were blocked for 30 min with 10% FBS in PBS-Tx, and incubated overnight at RT with rabbit anti-Wfs1 (1:500, Protein Tech, 11558-1-AP) in 1% FBS PBS-Tx. After three washes in PBS, sections were incubated for 2 h with donkey anti-rabbit Alexa Fluor488 (1:200, ThermoFisher Scientific, A-21206) in 1% FBS PBS-Tx, washed twice in PBS and mounted using ProLong Gold Antifade mountant (ThermoFisher Scientific, P10144).

Multichannel fluorescence stacks were achieved in a confocal microscope (Leica SP5) with LAS AF software v2.6.0 build 7266 (Leica), with a 40x objective (HCX PL APO CS 40.0×1.25 OIL UV), at 1 μm z-interval. Following lasers (excitation wavelength, emission spectral filter) were used for each fluorophore: Argon (488 nm, 499-535 nm) for Alexa Fluor488, DPSS (561nm, 571-620 nm) for Atto 550, and HeNe (633 nm, 652-738 nm) for Atto 647. Diode (405 nm, 415-485 nm) was used as an unstained channel to identify autofluorescent material, which can be abundant in epileptic tissue.

To estimate the amount of *Spag5* and *Dcc* transcripts, we calculated the number of dots per cell using ImageJ (Fiji). In one confocal plane per image, we counted single signal dots, in either Dcc or Spag5 channels, in each ROI. ROIs were drawn as the outline of several pyramidal cell somas, based on their Wfs1 immunoreactivity. Only those which were focus at their centre in the selected plane were drawn. When we found clusters instead of individual dots, we converted intensity to dot number as suggested by the manufacturer (ACD): (total intensity-average background intensity x area)/single dot average intensity.

### Statistical analysis

Statistical analysis was performed with MATLAB, SPPSS and using the computing environment R (R Development Core Team, 2005). No statistical method was used to predetermine sample sizes. Normality and homoscedasticity were evaluated with the Kolmogorov–Smirnov and Levene’s tests, respectively. The exact number of replications for each experiment is detailed in text and figures. Several ways ANOVAs or Kruskal-Wallis tests were applied. Post hoc comparisons were evaluated with the Tukey-Kramer, Student or Wilcoxon tests. Deep-superficial trends were evaluated using Spearman correlation and tested against 0 (i.e., no correlation was the null hypothesis) at p<0.05 (two sided).

## Data and code availability

Two open R Shiny applications provide visualization of LCM-RNAseq (http://lopezatalayalab.in.umh-csic.es/CA1_Sublayers_&_Epilepsy/) and snRNAseq (http://lopezatalayalab.in.umh-csic.es/CA1_SingleNuclei_&_Epilepsy/) data.

## Supplementary Materials

Supplementary tables are included in MS Office Excel format as follows:

Table S1: Electrophysiological properties of deep and superficial cells.

Table S2: LCM-RNAseq data.

Table S3: snRNAseq data.

**Fig.S1.**
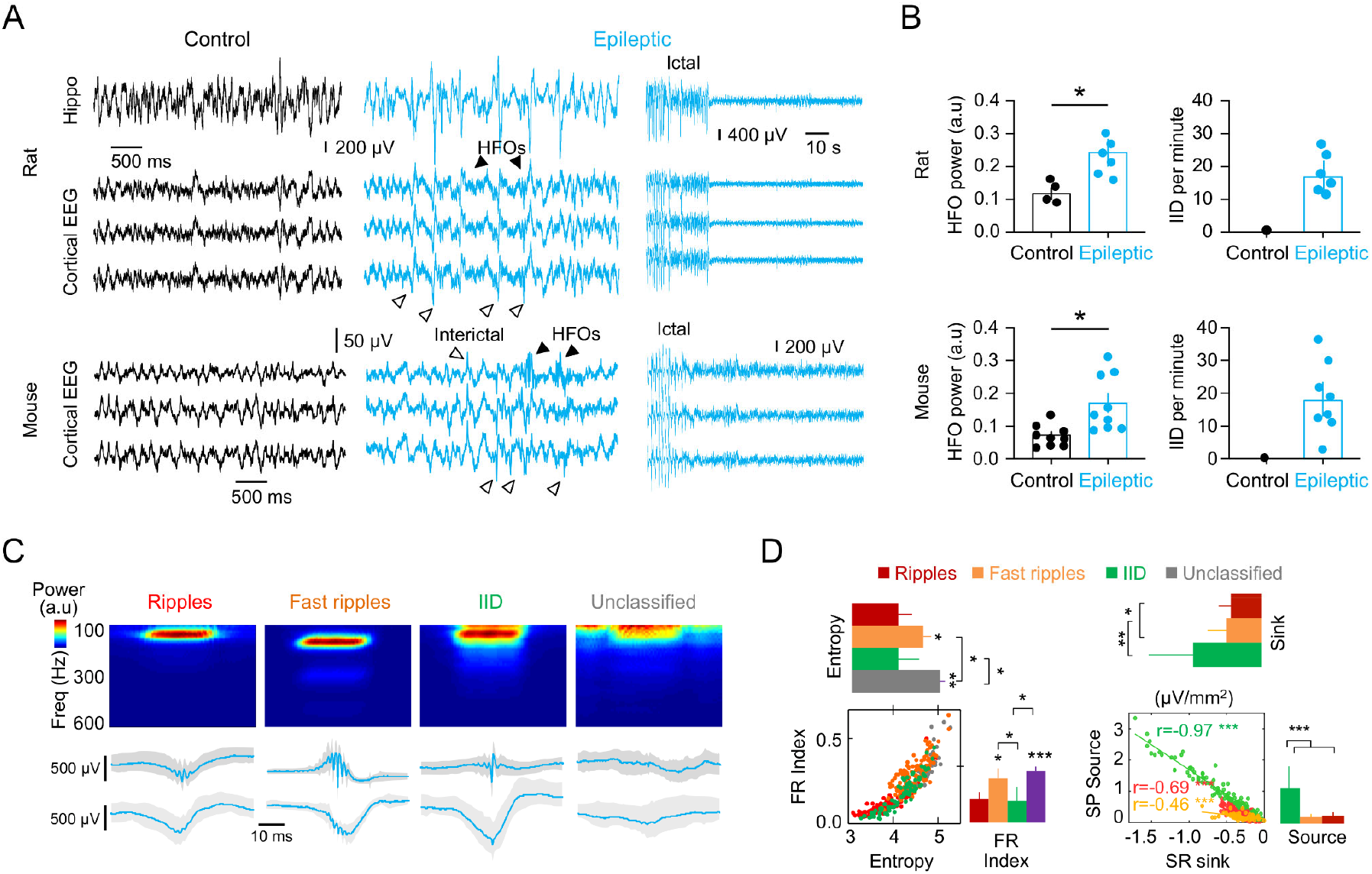
Electrophysiological analysis. **A**, Cortical EEG and/or intrahippocampal recordings from epileptic rats and mice allowed evaluation of neurophysiological changes accompanying acquired epilepsy. Simultaneous cortical and hippocampal recordings are shown for rats, while cortical EEG is shown for mice. Note interictal discharges (IID) and high-frequency oscillations (HFOs) in epileptic animals recorded both intracranially and at the cortical surface over the hippocampus. The small size of the mouse brain and connectivity between the hippocampus and prefrontal cortex facilitates recording of HFOs at scalp. The late stage of a seizure recorded in both animals is shown to highlight ictal activity. **B**, Quantification of the HFO power and IID rate in rats and mice 6 weeks after status. IID rate was evaluated from 20 min sessions in epileptic animals. Data from 1 session from 6 rats and 3 sessions from 3 mice. *, p<0.05 unpaired t-test. **C**, Sharp-wave (SPW) associated HFO events recorded from the dorsal hippocampus of urethane anesthetized epileptic rats were automatically detected and separated as ripples (100-150 Hz), fast ripples (>150 Hz) and IID using amplitude and spectral information. Some events were left unclassified. **D**, Quantitative separation of ripples, fast ripples and IID events. Spectral indices such as entropy and fast ripple indices were combined with information on the amplitude of current-source density (CSD) signals. IID were clearly separated from SPW events using CSD amplitude of sinks and sources. Asterisks reflect significant differences from post hoc Tuckey tests as *, p<0.05; **, p<0.01 and ***, p<0.001.

**Fig.S2.**
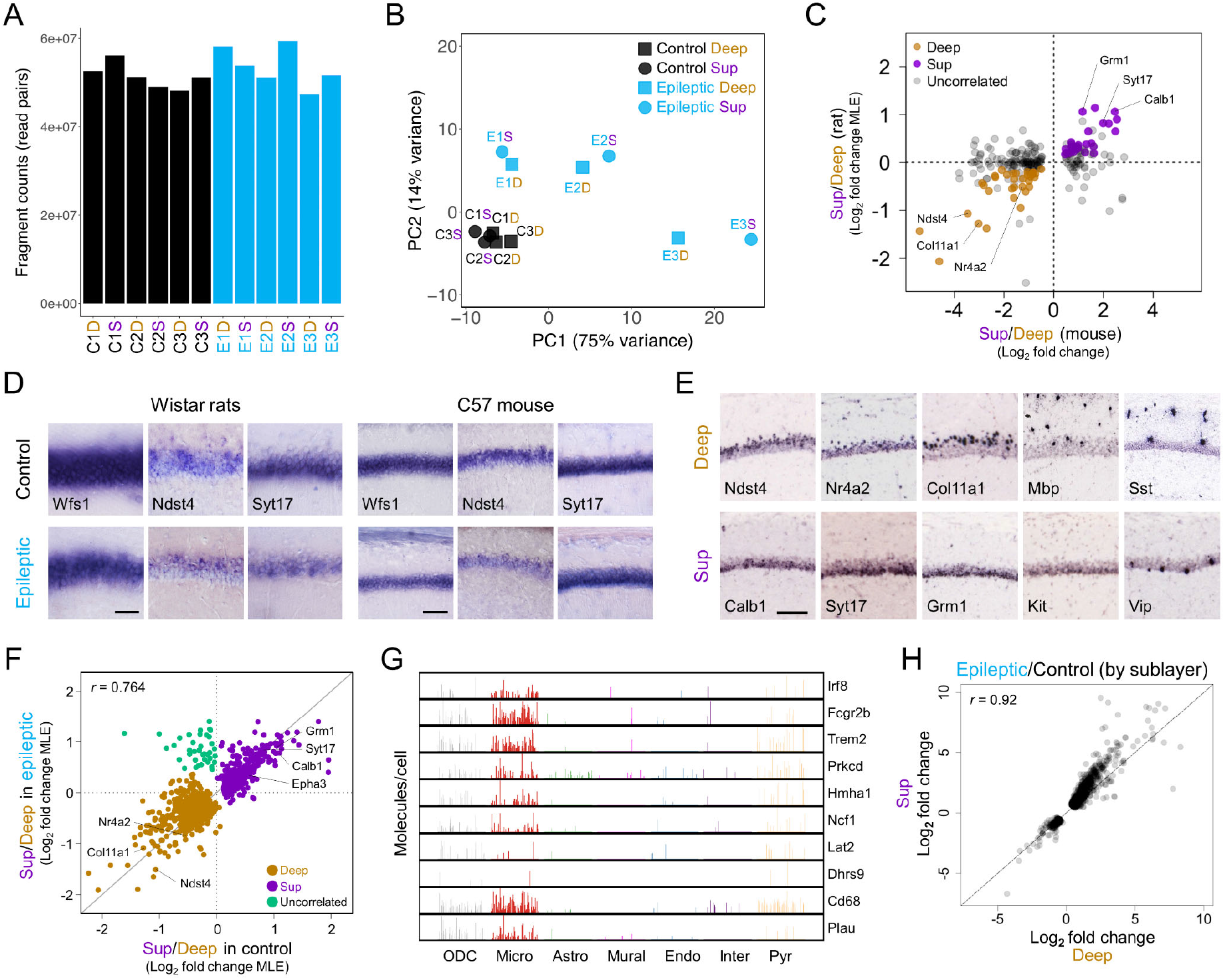
CA1 sublayer-specific gene expression profiling from bulk tissue (LCM-RNAseq). **A**, Bar graph showing per-sample sequencing depth. Note similar library size of about 50*10^6^ fragments per sample in both groups (paired-end sequencing). **B**, Principal component analysis of normalized rlog counts. Note pairwise clustering of deep and superficial samples from the same animals and significant separation between control and epileptic replicates, suggesting that transcriptional variability successfully captures between-group differences. **C**, Scatterplot showing differentially expressed genes (false discovery ratio, FDR<0.1) between superficial and deep manually sorted neurons from mice (x-axis; data from Cembrowski et al., 2016) against change in mRNA expression for these genes in our bulk-tissue LCM-RNAseq data from rats (y-axis). Colored points indicate genes that were differentially expressed across sublayers in both species. **D**, ISH of the pan-CA1-specific marker *Wfs1*, the superficial sublayer marker *Syt17* and deep sublayer marker *Ndst4* in representative hippocampal sections from control and epileptic rats and mice. Note sublayer markers are not all-or-none, but rather reflect a gradient distribution. Scale bar, 100 μm. **E**, Allen Brain Atlas (ABA) ISH sections showing additional genes differentially regulated in deep and superficial sublayers in mouse. Note regionalization of some interneuron-specific genes such as Sst and Vip, as well as non-neuronal genes such as Mbp. Scale bar, 100 μm. **F**, Scatter plot of DEGs in superficial and deep CA1 sublayers from control and epileptic rats (Adj p-val<0.05). Most transcripts are highly correlated, except for a subset of transcripts (green). **G**, Bar plot of the eleven cell-type-specific uncorrelated DEGs from F suggest contribution of different cell-type populations, especially microglia. **H**, Scatter plot showing changes in gene expression upon epilepsy in CA1 superficial and deep sublayer. Plot includes all differentially expressed genes in epilepsy (global effect: 732 unregulated, 185 downregulated genes (Adj p-val<0.01 and LFC>0.5).

**Fig.S3.**
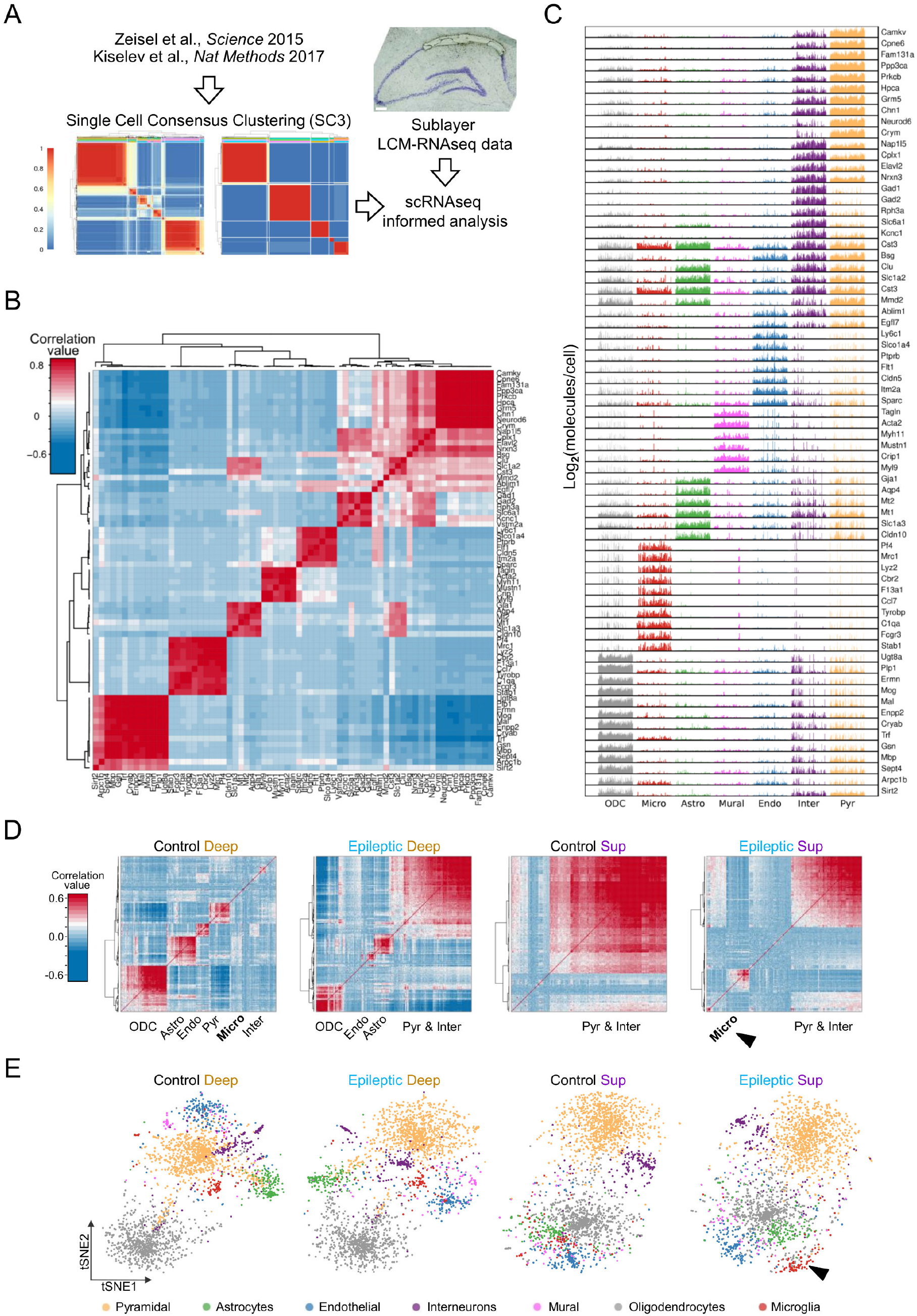
Single-cell informed analysis of LCM-RNAseq data from bulk hippocampal CA1 tissue. **A**, Workflow analytic steps to identify cell-type heterogeneity in CA1 hippocampal area from bulk tissue RNAseq data. Single-cell consensus clustering (SC3) analysis (Kiselev et al., 2017) was performed on previously published single-cell RNAseq data from mouse cortex and hippocampus (Zeisel et al., 2015) to identify major gene markers of cell types in CA1 area. Next, snRNAseq data for genes identified in LCM-RNAseq analyses (gene lists of top differentially expressed genes between CA1 sublayers of control and epileptic rats) was used to identify the presence of cell type gene markers and co-expressed genes at single cell level. **B**, A list of 69 cell-type gene markers was obtained using SC3 on single-cell RNAseq data from Zeisel et al. (2015). **C**, Bar plot showing expression levels as log2(molecules/cell) of the 69 cell type gene markers at single cell level (x-axis) across the seven major cell types identified. Pyr, pyramidal neurons; Inter, interneurons; ODC, oligodendrocytes; Astro, astrocytes; Endo, endothelial cells; Micro, microglia; Mural, mural cells. **D**, Clustered correlation matrices of expression levels of differentially expressed genes between deep and superficial CA1 sublayers from control and epileptic rats. Sets of differentially expressed genes of the same size (top 250 DEG) were used to investigate cell type heterogeneity in each condition. Note distinct cell-types segregate across sublayers. Note also larger number of microglial transcripts in response to epilepsy in superficial sublayer (arrowhead). **E**, t-Distributed Stochastic Neighbor Embedding (t-SNE) maps were generated using scRNAseq data for genes identified as differentially expressed between deep and superficial CA1 sublayers from control and epileptic rats. Sets of differentially expressed genes (top 250 DEG) were used to investigate cell type heterogeneity in each condition (max. Adj p-val = 0.029).

**Fig.S4.**
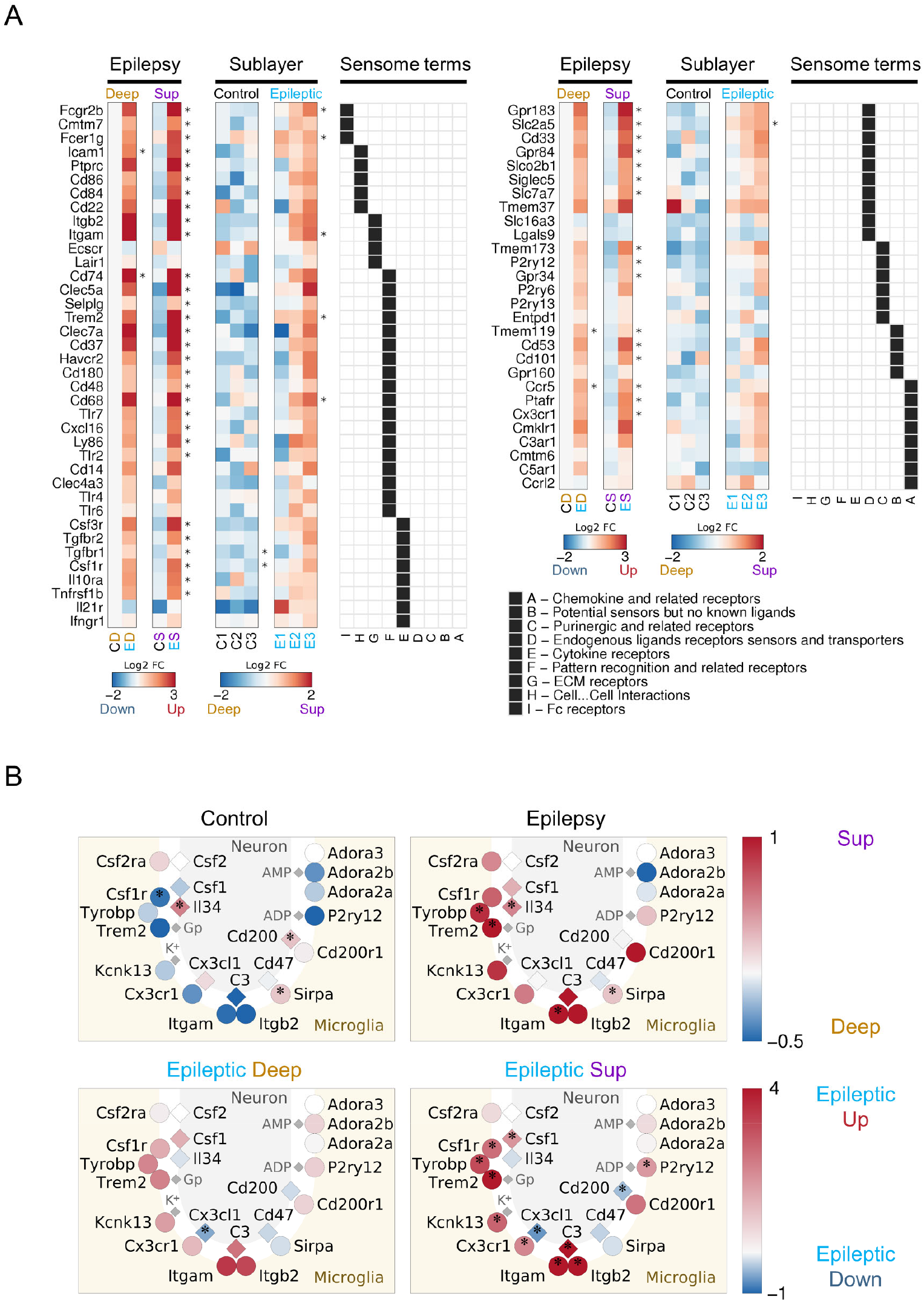
Transcriptional changes associated with microglia in the epileptic CA1 (bulk tissue LCM-RNAseq). **A**, Heatmaps of log2 fold changes for genes annotated to the microglial sensome (Hickman et al., 2013). Log2 fold change values are for the epilepsy effect in superficial and deep sublayers (Epilepsy), and for sublayer effect in control and epileptic mice (Sublayer). * Adj p-value<0.1. Gene Ontology annotation (GO BP) is shown at the foremost right (black squares). The list continues from left to right. **B**, Schematic diagram illustrating neuron-glia interactions via ligand-receptor pairings. Neuronal ligands (diamonds) and microglia receptors (circles) are colored by expression fold change (see color scheme legend) in the comparison between superficial and deep sublayer in control (top-left) and epileptic (top-right) CA1, and for epilepsy effect in deep (bottom-left) and superficial (bottom-right) CA1 sublayer. * Adj p-value<0.1.

**Fig.S5.**
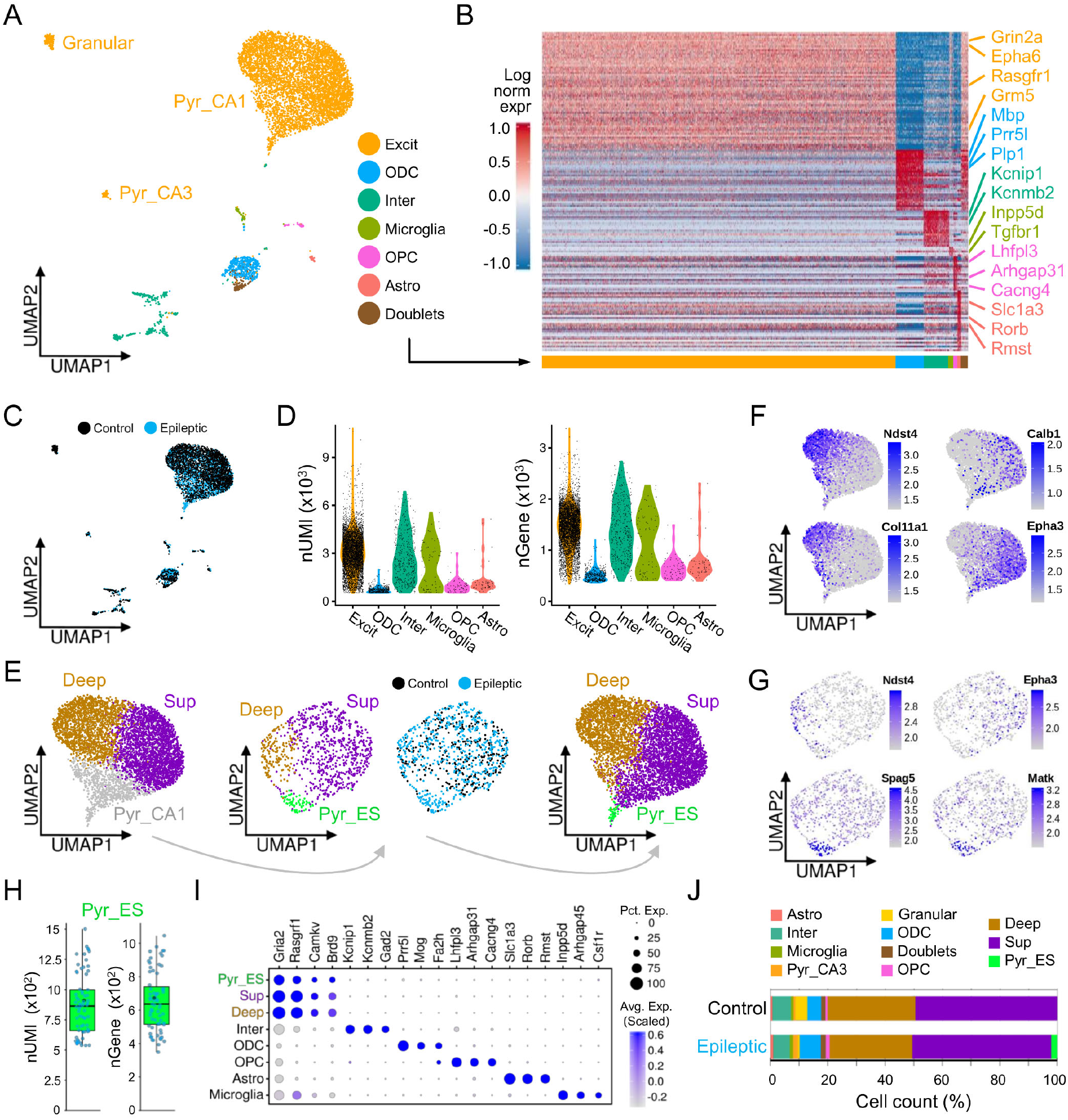
Transcriptional profiling at single-nucleus level unfolds heterogeneity of excitatory pyramidal neurons in the basal and epileptic hippocampal CA1 area. **A**, Uniform Manifold Approximation and Projection (UMAP) plot from snRNAseq transcriptional profiling showing 6 major cell classes identified and annotated in our CA1 samples. **B**, Heatmap of cell-type marker genes (168 enriched genes with AUC power>= 0.55) expression per single-nucleus across identified cell populations. **C**, UMAP plot of the snRNAseq datasets split by condition. **D**, Violin plots showing distribution of nUMI (left) and nGene (right) per major populations as indicated in the legend. **E**, UMAP clustering of the CA1 pyramidal cell population took two rounds. In the first round, nuclei were automatically separated in three main subclusters: deep cells, superficial cells and a third cluster of pyramidal neurons (Pyr_CA1). In the second round, the subsetted Pyr_CA1 cells were sorted as deep and superficial and epilepsy-specific Pyr_ES cells (green). **F**, UMAP of the pyramidal cell class colored by normalized expression levels for the indicated subpopulation gene markers. **G**, UMAP of the subsetted pyr_CA1 pyramidal cells. Note separation of deep and superficial cells, as well as the epilepsy-specific pyramidal cell subpopulation (Pyr_ES). **H**, Box plots showing nUMI (left) and nGene (right) per cell in the epilepsy-specific population Pyr_ES population. **I**, Confirmation of cell-type specific gene mapping of sorted cells. **J**, Proportions of the distinct cell types and populations identified across conditions (control and epilepsy). Note the epilepsy-specific population Pyr_ES (light green).

**Fig.S6.**
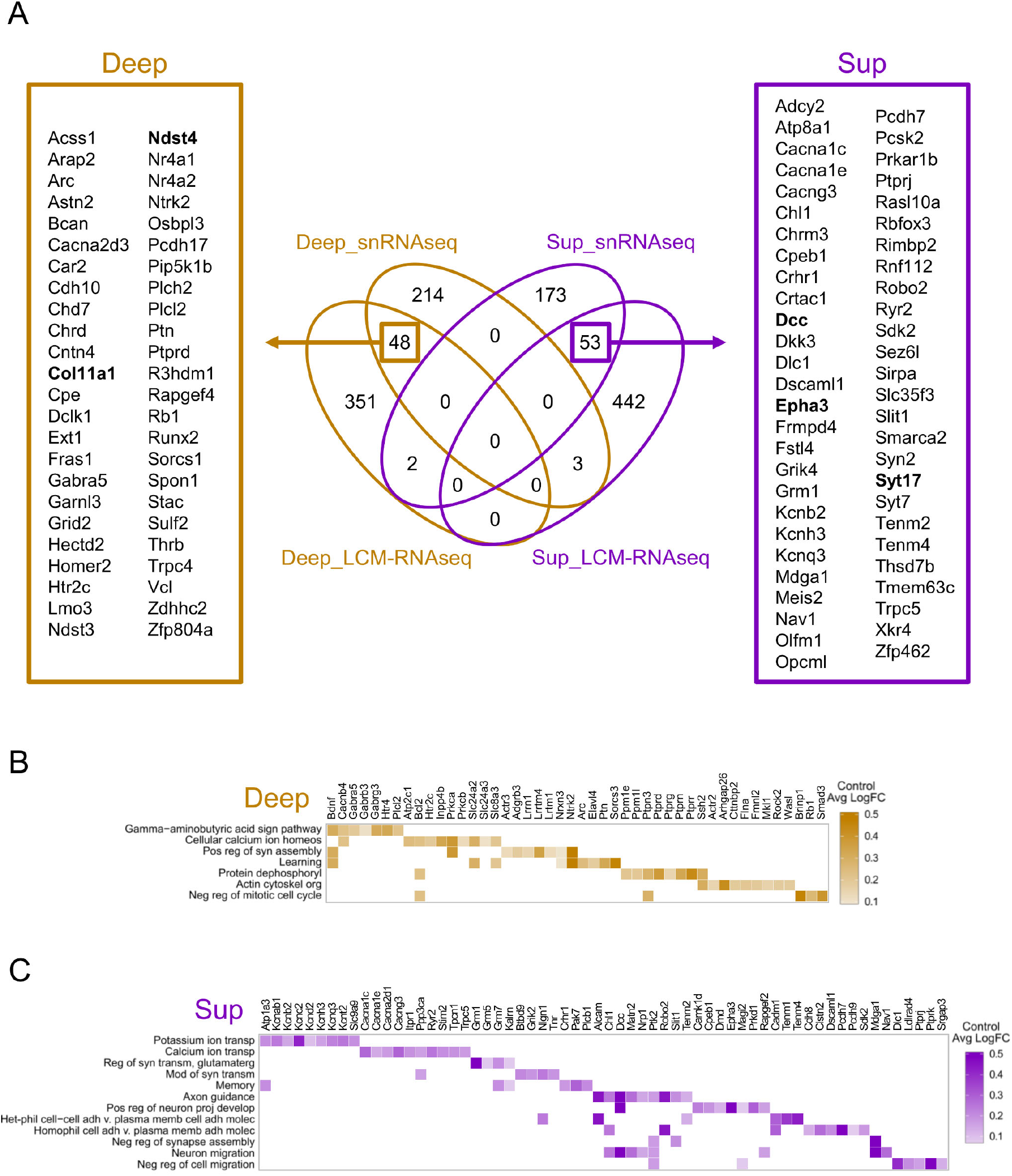
Equivalence between LCM-RNAseq and snRNAseq sublayer-specific results. **A**, Venn diagram showing overlap in genes identified as significantly enriched in superficial (Sup_LCM_RNAseq) and deep sublayer (Deep_LCM-RNAseq) by bulk CA1 sublayer-specific tissue gene expression profiling (LCM-RNAseq) or snRNAseq. Many sublayer- and cell-subtype significantly enriched transcripts were common across species (rat, mouse) and technologies (bulk RNAseq, snRNAseq). **B**, Functional GO analysis of genes significantly enriched in deep (top) or superficial (bottom) CA1 neurons (snRNAseq) in the control animals.

**Fig.S7.**
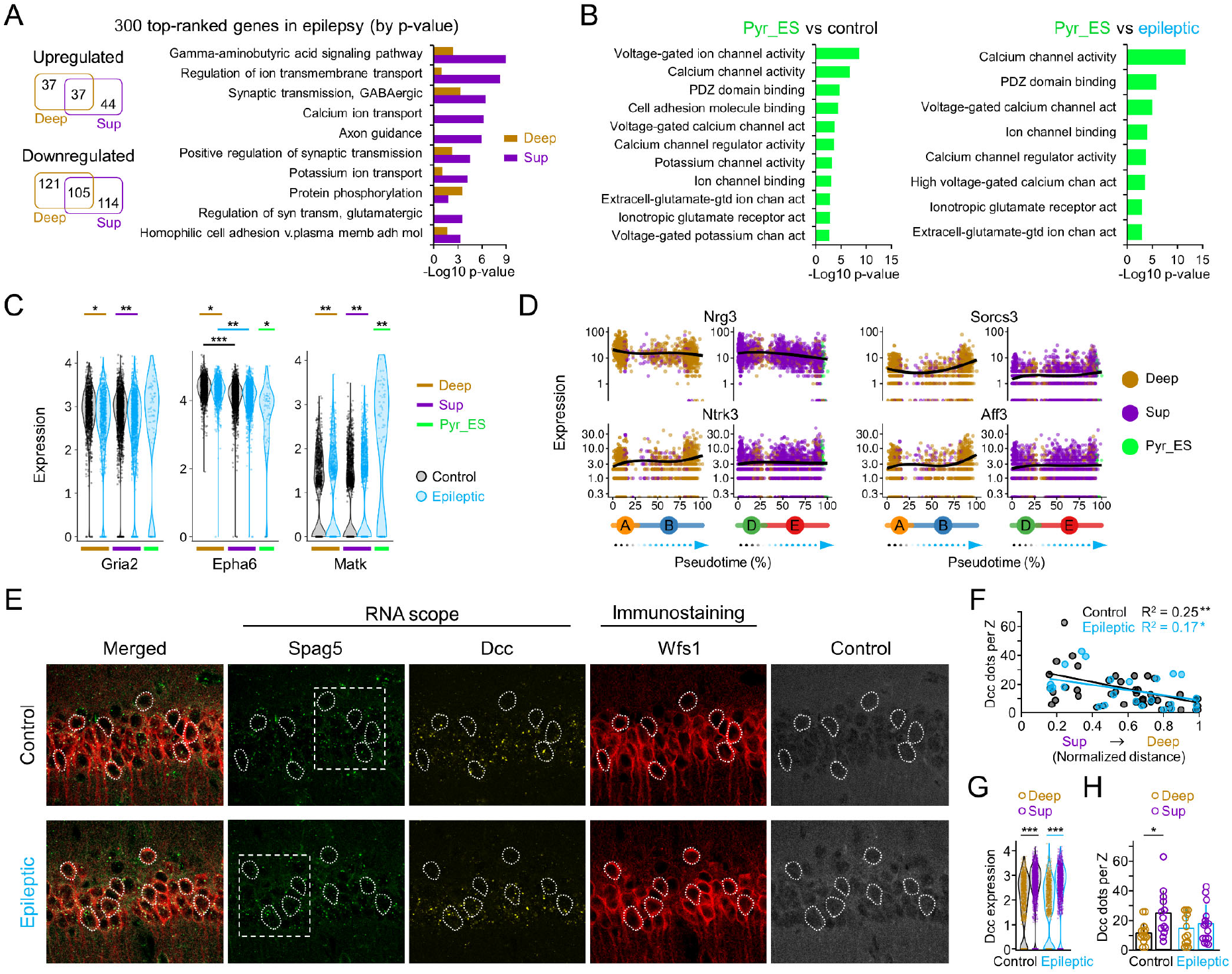
Gene profiling at single cell nuclei reveals transcriptional changes in superficial and deep CA1 neurons in experimental epilepsy. **A**, Venn diagram and Go terms of top-ranked 300 genes modulated by epilepsy. **B**, Bar chart of significance (left) and fold change (right) for most upregulated genes in Pyr_ES when compared with epileptic CA1 pyramidal cells (absolute fold change >0.5 and transcript detection in >50% of the nuclei). **C**, Violin plots showing normalized expression value (normalized log transformed UMIs) by condition (control, black; epilepsy, blue) and population (deep, occher; superficial, purple; Pyr_ES, lightgreen) of selected gene markers of pyramidal neurons (*Gria2, Epha6*), and *Knain2* which is modulated by epilepsy in superficial and deep cells. Note specific upregulation in Pyr_ES cells. *p<0.05; **p<1E-10; ***p<1E-50 (Wilcoxon rank sum test). **D**, Examples of gradient progression across disease trajectory for significantly modulated genes in epilepsy related to neuronal survival (*Nrg3, Ntrk3, Sorcs3, Aff3*). **E**, Combined RNAscope and immunostaining analysis allowed identification of cells with extreme expression of *Spag5* in situ. Neurons having their soma cut transversally by the confocal plane are outlined. Discontinuous line boxes identify the region expanded in Fig.5. The channel Control (405 nm) was used to control for autofluorescent signals across channels, which are characteristic of epileptic tissue (labeled Control at the rightmost). **F**, Distribution of *Dcc* dots per cells along their normalized position in the deep-superficial axis. Note similar trends for control and epileptic cells (significant Pearson correlation). **G**, Violin plot for *Dcc* gene (normalized log transformed UMIs) in nuclei of CA1 superficial (Sup) and deep (Deep) pyramidal cells from control (black) and epileptic (blue) animals. ***p<1E-50 (Wilcoxon rank sum test). **H**, Quantification of *Dcc* signals in deep and superficial CA1 pyramidal cells as counted in one confocal plane from 3 control and 3 epileptic mice (2-way ANOVA effect for sublayer F(1, 51)=6.771, p =0.012; no group differences nor interaction). Data from 27 control (14 deep, 14 superficial) and 28 epileptic cells (15 deep, 13 superficial).

## Acknowledgments

**General**: We thank Beatriz Lázaro, Ester Lara, Laura Dolón and Juan Moriano for technical help.

## Author contributions

LMP, JPLA, ABayes and ABarco designed the study. LMP and JPLA coordinated experiments and analysis. EC, MV, BG, DM, CMN, LBE, RRV, AVM, LD, IFL and DGD obtained data. YH and MS provided the Thy1.2-G-CaMP7-T2A-DsRed2 mouse line. EC, AMG, MV, BG, AB, JPLA and LMP analyzed and interpreted the data. AMG developed online tools. LMP and JPLA drafted the paper. All authors commented and contributed.

